# Predicting cochlear synaptopathy in mice with varying degrees of outer hair cell dysfunction using auditory evoked potentials

**DOI:** 10.1101/2025.05.09.653157

**Authors:** Brad N. Buran, Seán Elkins, Wenxuan He, Naomi F. Bramhall

## Abstract

Human temporal bones suggest a steady decline of cochlear synapses with age and greater synapse loss in adults with a history of military or occupational noise exposure. However, there is currently no validated method of diagnosing this type of cochlear deafferentation in living humans. Animal models indicate that cochlear synaptopathy is associated with reduced auditory brainstem response (ABR) wave 1 amplitude and envelope following response (EFR) magnitude for a sinusoidally amplitude modulated (SAM) tone. However, translating the SAM EFR to humans is complicated because it is difficult to obtain this measurement in humans using the same modulation frequency that showed the strongest relationship with synaptopathy in mice (1000 Hz). Computational modeling suggests that EFR magnitude measured with a rectangular amplitude modulated (RAM) tone may be a more sensitive measure of synaptopathy than the SAM EFR. In addition, because synaptopathy likely co-occurs with outer hair cell dysfunction, a diagnostic assay for synaptopathy needs to be robust even when auditory thresholds are abnormal. This study compared the relative ability of the ABR, SAM EFR, and RAM EFR to predict synapse numbers in mice with a large range of auditory thresholds and degrees of synaptopathy. The results indicate that the RAM EFR modulated at 1000 Hz is the single best predictor of synapse number when there is a broad loss of synapses across frequency, while combining RAM EFR and ABR further improves synapse prediction. In contrast, focal synaptopathy is best predicted by ABR wave 1 amplitude.

**Significance Statement:** This study assessed the relative ability of two auditory evoked potentials to identify cochlear synaptopathy, a type of cochlear deafferentation that occurs with age and noise exposure, in mice. Performance of these measures in the presence of outer hair cell (OHC) damage was also evaluated because synaptopathy is expected to often co-occur with OHC dysfunction. Concrete recommendations of measurements to use for non-invasive diagnosis of synaptopathy in humans are provided. This represents a significant advance toward diagnosis of a condition that is thought to have a high prevalence in humans. The ability to identify individuals with cochlear synaptopathy is vital for furthering our understanding of how this auditory deficit impairs auditory perception and the future development of treatment options.

## Introduction

Cochlear synaptopathy, which is the loss of the synapses between the inner hair cells (IHCs) and the afferent auditory nerve fibers, results from aging and noise exposure in animal models (Kujawa and Liberman, 2009; Sergeyenko et al., 2013). Human temporal bones suggest that synaptopathy also occurs in humans with age (Wu et al., 2019) and military/occupational noise exposure (Wu et al., 2021). Due to the high predicted prevalence of synaptopathy, the development of a diagnostic measure is of high importance.

Although synaptopathy can only be confirmed through post-mortem analysis, two auditory evoked potentials have been identified in mice as possible non-invasive indicators of synaptopathy; the auditory brainstem response (ABR) wave 1 amplitude (Kujawa and Liberman, 2009; Sergeyenko et al., 2013) and the envelope following response (EFR; Parthasarathy and Kujawa, 2018; Shaheen et al., 2015).

ABR wave 1 amplitude may underestimate synaptopathy because it is an onset response that is unaffected by the loss of low spontaneous rate auditory nerve fibers (Bourien et al., 2014), a subset of fibers that appears to be particularly vulnerable to synaptopathy (Furman et al., 2013; Schmiedt et al., 1996). The EFR measures the sustained response to an auditory stimulus, rather than the onset response. However, translating the EFR from mouse to humans presents challenges. Higher modulation frequencies (f_m_) are thought to reflect auditory nerve function (Parthasarathy and Kujawa, 2018; Shaheen et al., 2015) while lower modulation frequencies appear to be generated primarily by the brainstem (Kuwada et al., 2002). In mice, the EFR is most sensitive to synaptopathy when f_m_=∼1000 Hz (Shaheen et al., 2015). While most studies of human synaptopathy use low modulation frequencies (f_m_=80-120 Hz) because high frequencies result in very small EFR magnitudes (Purcell et al., 2004), McHaney et al. (2024) showed that it is possible to record EFRs in young adults at f_m_=1024 Hz. Although the EFR has only been measured in animal models of noise-or age-related synaptopathy using a sinusoidally amplitude modulated (SAM) stimulus, computational modeling suggests that a rectangular amplitude modulated (RAM) stimulus may be more sensitive to synaptopathy due to its sharply rising envelope (Vasilkov et al., 2021). In budgerigars with kainic acid-induced synaptopathy, Garrett et al. (2025) showed a greater reduction in EFR magnitude compared to controls using a RAM versus a SAM stimulus (f_m_=100 Hz). Adding further to uncertainty about how to use the EFR to detect synaptopathy, the methods for calculating EFR strength have varied across human studies (e.g., Van Der Biest et al., 2023; Zhu et al., 2013).

Two studies obtained ABR/EFR measures and synapse numbers in the same animals. Shaheen et al. (2015) compared ABR and SAM EFR (f_m_=1000 Hz) in mice with noise-induced synaptopathy and reported larger effect sizes for SAM EFR. Parthasarathy and Kujawa (2018) evaluated ABR and SAM EFR (f_m_=1028 Hz) in mice with age-related synaptopathy and observed larger effect sizes for the EFR at 30 kHz, but the opposite at 12 kHz. This provides modest support for the SAM EFR (f_m_=∼1000 Hz). However, it’s unclear how the RAM EFR compares to the SAM EFR and ABR. The best measure may depend on the synaptopathy configuration. In mice, noise-induced synaptopathy initially results in a focal loss of synapses from 22.6-64 kHz (Kujawa and Liberman, 2009) which broadens over time to include lower frequencies (Fernandez et al., 2015). Broad synapse loss also occurs in mice with age-related synaptopathy (Sergeyenko et al., 2013) and in older humans (Wu et al., 2019). Another important consideration is how each measure is impacted by OHC dysfunction, which likely co-exists with synaptopathy in many cases.

The objective of this study was to evaluate the relative ability of evoked potential measures to predict synapse numbers in mice with varying degrees of synaptopathy and OHC dysfunction. The measures included ABR wave 1 amplitude, SAM EFR (f_m_=110 Hz and 1000 Hz), and RAM EFR (f_m_=110 Hz and 1000 Hz). Different methods of EFR processing were also investigated.

## Materials and Methods

### Experimental Design

Four groups of CBA/CaJ mice (Jackson Laboratories #000654, https://www.jax.org/strain/000654) were included in this study: 1) young (n=17), 2) acute noise exposed (n=13), 3) aged (n=14), and 4) aged after noise exposure (n=13). The goal was to generate a sample of mice that encompassed a wide variety of degrees of cochlear synapse loss with and without varying degrees of OHC damage. The young mice were tested between 10 and 42 weeks of age. The acute noise exposed mice were exposed to 94 or 98 dB SPL at 8 weeks of age and tested between 10 and 13 weeks of age. The aged mice were tested between 82 and 103 weeks of age. The aged after noise exposure mice were noise exposed to 101 dB SPL at 16 weeks of age and tested between 80 and 104 weeks of age. Both the acute noise exposed and aged after noise exposure groups were exposed to octave-band (8 to 16 kHz) noise for two hours. All physiological and histological measures were performed in the left ear for each mouse. Mouse data were included in the dataset even if a full set of physiological and histological measures were not available (e.g., ∼33% of experiments did not measure the SAM and RAM EFR modulated at 1 kHz for a carrier frequency of 32 kHz and ∼30% of experiments did not have a measurable ABR wave 1 amplitude at 40 dB SPL). Mice of both sexes were included.

### Noise exposure

Noise exposures were performed following Kujawa and Liberman (2009). Awake mice were placed in a custom-built wire cage inside a custom-built noise exposure chamber. The cage was subdivided into six compartments with one mouse per compartment. The cage was positioned on a turntable, ensuring that all mice received a uniform noise exposure that was not affected by the development of standing waves in the chamber. The noise exposure chamber was designed as a trapezoidal prism with only the top and bottom surfaces parallel to each other. Noise exposure was controlled using a custom-written program (Buran, 2024c). Noise stimuli were generated digitally by drawing random samples from a uniform distribution and then bandpass filtered using a 1001 tap finite impulse response filter with a passband of 8 to 16 kHz (> 60 dB/octave slope). To ensure uniform spectral distribution of the noise, the gain of the passband was adjusted to equalize the noise. Stimuli were converted to analog (PCI-6251, National Instruments), amplified (D75-A, Crown Audio), and delivered via a 1-inch compression driver (D220 Ti, JBL Professional Loudspeakers) coupled to a horn (HM25-25, JBL Professional Loudspeakers). Prior to each noise exposure, the intensity level of the noise was calibrated to the target level using a ¼-inch microphone (377C01 coupled to a 426B03 preamplifier and powered by a 480C02 signal conditioner, PCB Piezotronics).

### Physiological measures

Animals were anesthetized with a ketamine (100 mg/kg) and xylazine (10 mg/kg) cocktail. Body temperature was monitored rectally and maintained using a heating pad regulated by a homeothermic temperature controller (50-7503F, Harvard Apparatus). A custom-built acoustic system consisting of two speakers and an embedded microphone (Buran et al., 2020) was positioned inside the intratragal notch just above the external acoustic pore (i.e., the opening of the ear canal). Stimuli were generated digitally using a custom data acquisition program (Buran and David, 2025), converted to analog (PXI-4461, National Instruments), amplified (SA-1, Tucker-Davis Technologies), high-pass filtered at 500 Hz using a custom-built RC circuit, and delivered to the ear via one of the speakers (CDMG15008-03A, Same Sky [formerly CUI Devices]) in the acoustic system. The embedded microphone (FG-23329-P07, Knowles Electret) was calibrated using a 1/8-inch microphone (46-DP1 powered by a 12AK, GRAS Acoustics). In-ear calibration of the speakers was performed immediately prior to each experiment using the embedded microphone. Responses to ABR and EFR stimuli were collected using needle electrodes (F-E2-12, Natus Medical) positioned at the vertex and intratragal notch with a ground near the tail. Responses were amplified (50,000x), band-pass filtered from 10-10,000 Hz (P511, Grass Instruments), and digitized (PXI-4461, National Instruments) for further analysis.

ABR stimuli consisted of 5 msec tone pips (0.5 msec cosine-squared rise-fall ramp with 4 msec steady-state) presented at a rate of 81/s. A full stimulus train consisting of a single presentation of each of seven frequencies (5.6 to 45.2 kHz in half-octave steps) and levels (10 to 80 dB SPL in 5 dB steps) was constructed. ABRs were presented in alternating polarity and a total of 512 artifact-free trials were collected (256 for each polarity). The ordering of frequencies and levels was arranged in the interleaved ramp design described in Buran et al. (2020) which leverages auditory nerve fiber tuning to minimize adaptation. Ensemble averages of ABR waveforms were bandpass filtered at 300 to 3000 Hz and the peak for wave 1 initially assigned using semi-automated peak-picking software (Buran, 2024a) and reviewed visually by an experienced rater. ABR thresholds were identified using an automated algorithm (Shaheen et al., 2025).

EFR stimuli consisted of 500 msec amplitude-modulated carrier tones with an inter-stimulus interval jittered uniformly between 100 and 120 msec for an average stimulus rate of 1.64/s. Carrier frequencies were 16 or 32 kHz with an overall stimulus level of 70 dB SPL. Stimuli were either sinusoidally (SAM) or rectangular amplitude modulated (RAM) at 110 or 1000 Hz. For SAM tones, the modulation depth was 100%. For RAM tones, the modulation depth was 100% and the duty cycle was 25%. A 2.5% Tukey window was applied to the onset and offset of each individual cycle of the RAM tone. Stimuli were presented in alternating polarity and a total of 128 trials were collected (64 for each polarity). As illustrated in **Figure 1**, five different methods of analyzing the EFR data were evaluated: 1) Compute the power at the modulation frequency (f_0_); 2) Sum the power of the first five multiples of the modulation frequency (f_0-4_); 3) Compute the power at the modulation frequency and subtract the noise floor (f_0_ re NF); 4) Sum the power of the first five multiples of the modulation frequency after subtracting the noise floor at each frequency (f_0-4_ re NF); and 5) Compute the phase locking value (PLV). For all methods, EFR magnitude was calculated similarly to the bootstrapping approach described in Zhu et al. (2013). In this approach, 128 trials were drawn with replacement, averaged, and the power spectrum was computed. Random draws were balanced across positive and inverted polarities (i.e., 64 trials from each polarity). This process was repeated 100 times to generate a distribution of the power for each frequency bin. The average value of the distribution at the modulation frequency and the first four harmonics was used to estimate the raw EFR response. To estimate the noise floor, the power in the third to seventh discrete Fourier transform (DFT) bin on either side of the frequency of interest was averaged, for a total of eight bins.

**Figure 1:**
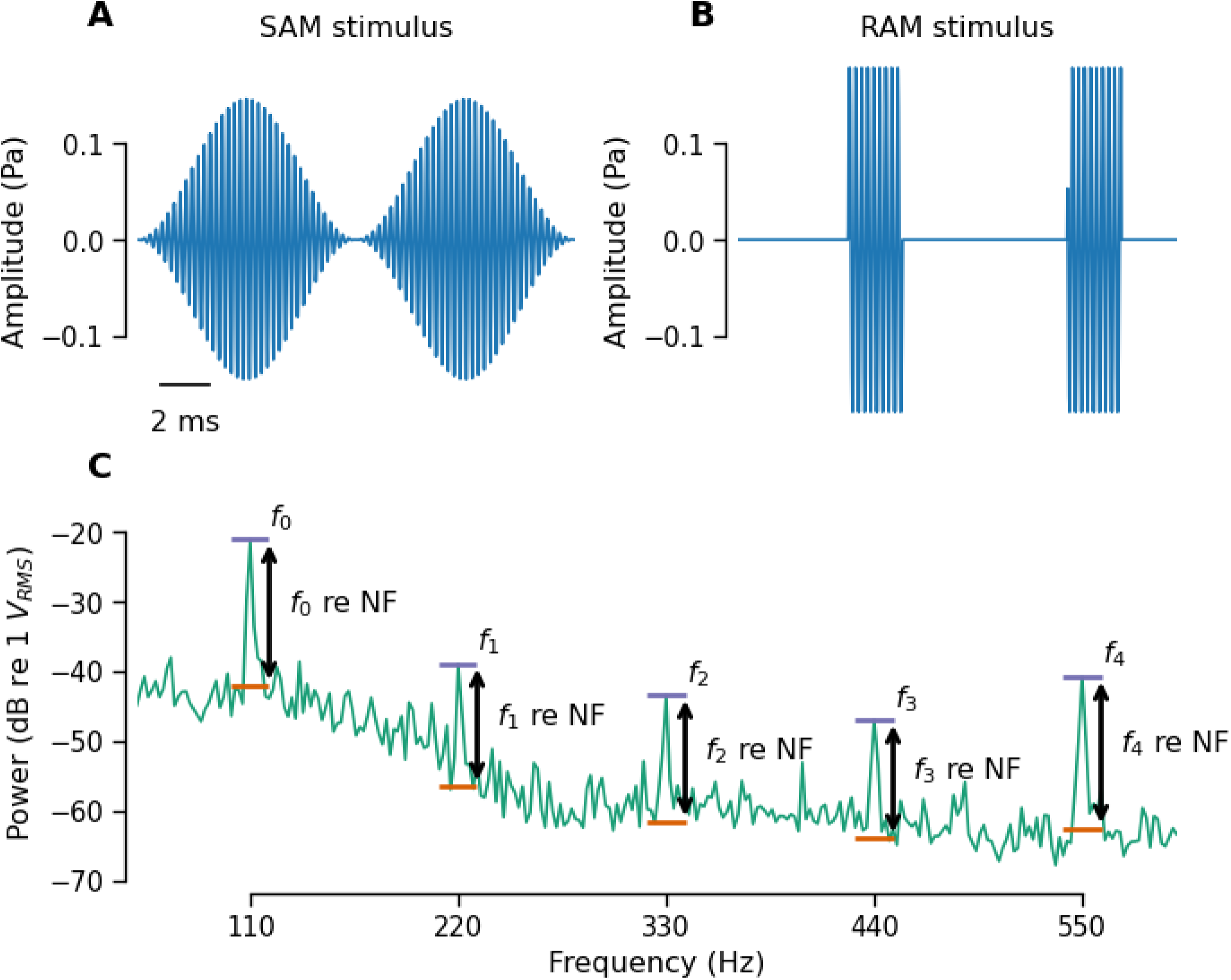
Schematic showing EFR stimuli and methods for analyzing the EFR response. Waveforms for SAM **(A)** and RAM **(B)** stimuli modulated at 110 Hz. Stimuli are scaled so that the overall RMS for both stimuli is 70 dB SPL. **(C)** Power spectral density of the EFR response to a RAM stimulus modulated at 110 Hz illustrating the features that are used in different methods of calculating EFR magnitude, including the noise floor (NF), the modulation frequency (f_0_) and the first five multiples of the modulation frequency (f_0_-f_4_). EFR power can either be extracted at f_0_ only or by summing f_0-4_ and can be expressed either as an absolute value (f_0_, f_0-4_) or relative to the noise floor (f_0_ re NF, f_0-4_ re NF). When summing f_0-4_, values are converted to a linear scale prior to summing and then converted back to a dB scale for analysis.

As an indicator of OHC function, distortion product otoacoustic emissions (DPOAEs) were recorded using the embedded microphone in the acoustic system. For all DPOAE measurements, the *f_1_* level (*L_1_*) was fixed at 10 dB higher than the *f_2_* level (*L_2_*) and the *f_2_*/*f_1_* ratio was fixed at 1.2. DPOAE thresholds were assessed using input-output functions at seven frequencies (5.6 to 45.2 kHz in half-octave steps). For each frequency, the *f_2_* level was swept from 10 to 80 dB SPL in 5 dB steps. Threshold was defined as the *f_2_* level at which the DPOAE level was 0 dB SPL. To parallel the DPOAE measurements often used in human studies of synaptopathy, DP-grams were obtained by sweeping the *f_2_* frequency in 1/8-octave steps from 5.6 to 45.2 kHz at two *f_2_* levels, 40 and 55 dB SPL.

### Histology

Immediately following the physiological measurements, mice were deeply anesthetized, decapitated, and the cochleae extracted for histology. A small hole was made in the apex of the cochleae and 4% paraformaldehyde in phosphate-buffered saline (PBS) at pH 7.3 was perfused through the round window. Cochleae were post-fixed for two hours at room temperature and then decalcified in 10% EDTA for 3-4 days. Once sufficiently decalcified, the cochleae were dissected into five pieces for whole mount immunostaining. Prior to immunostaining, the dissected pieces were cryoprotected in 30% sucrose and then permeabilized using a freeze-thaw step. If immunolabeling could not be initiated immediately, the cochlear pieces were stored at −80°C after the initial freezing step. Once thawed, cochlear pieces were rinsed in PBS, blocked with 5% normal horse serum (NHS) in PBS for one hour with 0.3% Triton-X added to further permeabilize the tissue. Primary antibodies were diluted in a solution of 1% NHS in PBS with 0.3% Triton-X. Cochlear pieces were incubated overnight at 37°C with rabbit anti-MyosinVIIa (dilution of 1:1000, Proteus Biosciences Cat# 25-6790, RRID:AB_10015251), mouse IgG1 anti-CtBP2 (dilution of 1:200, BD Biosciences Cat# 612044, RRID:AB_399431), and mouse IgG2a Rabbit anti-GluR2 (dilution of 1:2000, Millipore Cat# MAB397, RRID:AB_2113875). Following a wash in PBS, cochleae were incubated for one hour at 37°C with goat anti-mouse IgG1 AF568 (Thermo Fisher Scientific Cat# A-21124, RRID:AB_2535766), goat anti-mouse Ig2a AF488 (Thermo Fisher Scientific Cat# A-21131, RRID:AB_2535771), and goat anti-rabbit AF647 (Thermo Fisher Scientific Cat# A-21245, RRID:AB_2535813). All secondary antibodies were used at a dilution of 1:1000 in 1% NHS plus 0.3% Triton-X. Following a wash in PBS, the signal was amplified by a second incubation in a freshly-made batch of the same set of secondary antibodies for one hour at 37°C. Pieces were washed in PBS and then labeled with DAPI (Invitrogen D1306) at a dilution of 1:5000. A final wash in PBS was applied prior to mounting and coverslipping the pieces using ProLong Diamond Antifade Mountant (Thermo Fisher P36965).

A cochlear frequency map was computed using a custom program (Buran, 2024b) that translates distance along the cochlear partition into frequency using the Greenwood function for mouse (Muller et al., 2005). Confocal z-stacks were acquired for each ear at seven frequencies (5.6 to 45.2 kHz in half-octave steps) using a 1.4 NA 63x oil-immersion objective on either a Zeiss LSM 980 or a Leica TCS SP5 confocal.

CtBP2 puncta were identified using Imaris (Oxford Instruments). Assisted by custom software, identified CtBP2 puncta were inspected to identify all CtBP2 puncta paired to a closely apposed glutamate receptor patch (GluR2). Curation of CtBP2-GluR2 pairs were performed only for 16 and 32 kHz. The synaptograms (synapse counts across cochlear frequency) shown in **Figure 2E** are based on the number of CtBP2-positive puncta (i.e., synaptic ribbons) whereas all other figures show the number of CtBP2-GluR2 pairs (i.e., synapses).

**Figure 2:**
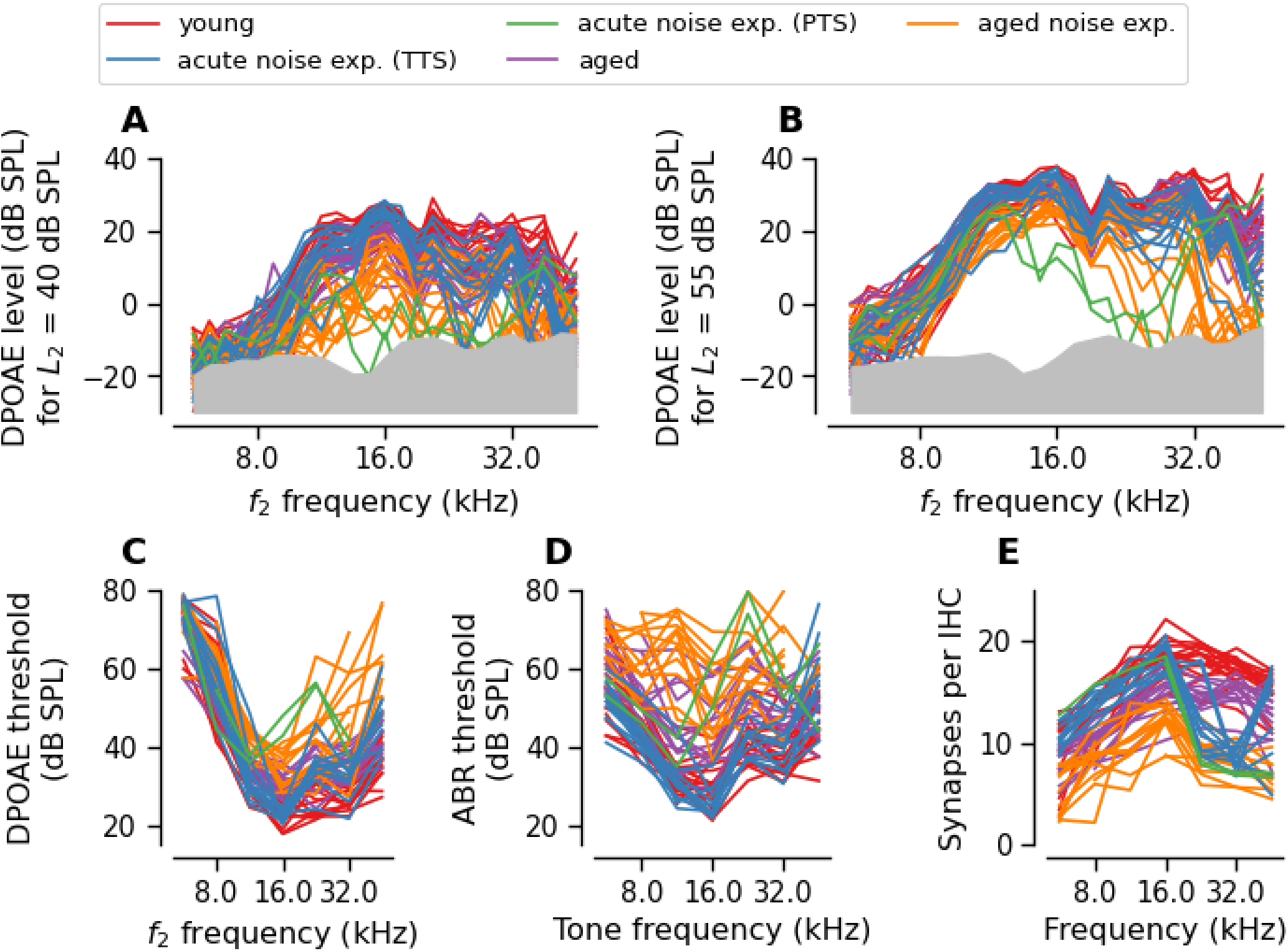
The sample included a large range of OHC dysfunction and synaptic loss. Plots show variation across individual mice from each experimental group for **(A-B)** DPOAE levels measured in a frequency sweep for an *f*_2_ fixed at 40 dB SPL **(A)** or 55 dB SPL **(B)**, **(C)** DPOAE thresholds, **(D)** ABR thresholds, or **(E)** the number of synapses per IHC across cochlear frequency (synaptogram). The acute noise exposed group was subdivided based on whether the mice experienced a temporary or permanent threshold shift (TTS or PTS). For all subsequent analyses, only data from 16 kHz and 32 kHz were used. In A and B, the shaded gray region indicates the DPOAE noise floor averaged across all ears in the sample.

All procedures were approved by OHSU’s Institutional Animal Care and Use Committee and conducted in accordance with guidelines from the Office of Laboratory Animal Welfare at the National Institutes of Health.

### Statistical Analysis

Due to time constraints imposed by the anesthesia, EFR data was only collected for 16 and 32 kHz carriers. Thus, the analysis of the physiological and histological data only include data from frequencies of 16 and 32 kHz. Data was collected only from the left ear of each mouse to minimize test session duration. Correlations between each of the measures were assessed using the Pearson correlation coefficient from the Python scipy library (Virtanen et al., 2020) and confidence intervals computed using the Fisher transformation.

To test the relative ability of each evoked potential measure to predict synapse numbers, we constructed a linear regression model based on various combinations of the evoked potential and DPOAE measures, 𝑦_𝑓,𝑖_ = 𝛽_0_ + 𝛽_1_ ⋅ Μ_𝑓,𝑖_ + 𝛽_2_ ⋅ 𝜉_𝑓,𝑖_ + 𝜖_𝑓,𝑖_.

The number of synapses for frequency 𝑓 in ear 𝑖 is represented by 𝑦_𝑓,𝑖_. The frequency and ear-specific evoked potential measure is represented by Μ_𝑓.𝑖_, the frequency and ear-specific DPOAE measure is represented by 𝜉_𝑓,𝑖_, and the residual error term is represented by 𝜖_𝑓,𝑖_. When testing 𝑝 combinations of evoked potential measures (e.g., ABR and EFR), the model was expanded to 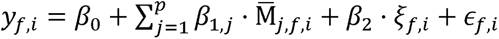. In this equation, evoked potential measure 𝑗 of frequency 𝑓 in ear 𝑖 is represented by Μ̅_𝑗,𝑓,𝑖_. Models were fit using the Python statsmodels library (Seabold and Perktold, 2010).

Model performance was assessed using ten repeats of 10-fold cross-validation (Burman, 1989). Data was partitioned into folds such that all observations from a single ear were assigned to the same fold. The acute noise exposed group of mice were not used for the model training since these mice had focal synaptopathy. Two sets of data were used for model validation: 1) the held-out subset from each fold of the cross validation and 2) a separate set of data from the acute noise exposed mice. Prediction error was quantified as the root-mean-squared error (RMSE), 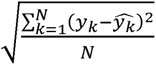, where 𝑦*_k_* is the k-th synapse (regardless of frequency) and 𝑦^_𝑘_ is the corresponding prediction. On each repeat, the data was shuffled such that the series of 10 folds contained different ears.

The initial models evaluated a total of six different evoked potential options (no evoked potentials, ABR wave 1 amplitude at 80 dB SPL [ABR_80_], SAM EFR modulated at 110 Hz [SAM_110_], SAM EFR modulated at 1000 Hz [SAM_1000_], RAM EFR modulated at 110 Hz [RAM_110_], and RAM EFR modulated at 1000 Hz [RAM_1000_]) and four approaches for adjusting for OHC function (no adjustment, DPOAE threshold, DPOAE level at an *L_2_* of 40 dB SPL [DPOAE_40_], and DPOAE level at an *L_2_* of 55 dB SPL [DPOAE_55_]).

To establish a baseline for interpreting the prediction error, an intercept-only model (i.e., 𝑦_𝑓,𝑖_ = 𝛽_0_ + 𝜖_𝑓,𝑖_) was tested in which the predicted synapse number is the average of the observed synapse numbers in the dataset. In addition, because OHC dysfunction is highly correlated with manipulations that reduce synapse numbers (i.e., aging and noise exposure), the ability of DPOAE measures to predict synapse numbers was also evaluated (i.e., 𝑦_𝑓,𝑖_ = 𝛽_0_ + 𝛽_2_ ⋅ 𝜉_𝑓,𝑖_ + 𝜖_𝑓,𝑖_). An ideal metric of synaptopathy would perform better at predicting synapse numbers than the intercept-only and the DPOAE-only models.

As an additional test of model performance, we refit each of these models without cross validation and computed the Akaike information criterion (AIC), which provides an estimate of the relative expected prediction error of different models after penalizing for the number of parameters. For this approach, the number of observations should be held constant across all models, so all combinations of ear and frequency for which we did not have a full set of observations were dropped, leaving a total of 83 observations from 52 unique ears (61 observations from 39 unique ears when excluding the acute noise exposed group). Since the sample size can be considered small relative to the model degree of freedom, we used a modification of the AIC that corrects for the increased risk of overfitting with small samples (AICc; Hurvich and Tsai, 1989). For model comparison, the difference in AICc between each model and the best-performing model was calculated (ΔAICc). While there is no hard rule of thumb, models with ΔAICc values ≤ 2 are generally considered comparable to the best performing model whereas models with ΔAICc values > 10 indicate a clear preference for the model with a lower AICc. Models with ΔAICc between 2 and 10 should not be rejected outright even though the evidence suggests that the model with a lower AICc is better.

### Code Accessibility

All custom-written software used for stimulus generation, data acquisition, and analysis are available on GitHub under permissive licenses (MIT, BSD 3-clause, etc.). See citations for details.

## Results

### Wide ranges of OHC function and synapse numbers were represented by the sample

Consistent with the goal of sampling mice with a heterogenous range of OHC function; DPOAE levels, DPOAE thresholds, and ABR thresholds spanned a large range of levels (**Figure 2A-D**, **Table 1**). Likewise, there was a broad range of synapse numbers across cochlear frequency in the sample (**Figure 2E**). Compared to the young mice, the aged mice and the mice who were aged after noise exposure had broad synaptopathy across the entire cochlea. The degree of synaptopathy increased with age and was accelerated by early noise exposure. In contrast, the acute noise exposed mice had focal synaptopathy with synapse numbers at 16 kHz and below comparable to the young mice, while synapse numbers at 32 kHz were reduced ∼50% relative to the young mice.

**Table 1.** Span of ABR threshold and DPOAE level/threshold measurements across all mice in the sample.

| Measure | 16 kHz span (dB) | 32 kHz span (dB) |
| --- | --- | --- |
| DPOAE level ($L_2 = 40$ dB SPL) | 39 | 44 |
| DPOAE level ($L_2 = 55$ dB SPL) | 25 | 43 |
| DPOAE threshold | 30 | 48 |
| ABR threshold | 50 | 55 |

### Relationship between OHC dysfunction and ABR/EFR measurements

Given that the ABR and EFR represent the summed activity of multiple auditory nuclei, a reduction in cochlear gain due to OHC dysfunction could alter the magnitude of these evoked measures. To evaluate the impacts of OHC dysfunction on the ABR and EFR, correlations between DPOAE level and either ABR wave 1 amplitude or EFR magnitude were examined at matching frequencies (16 kHz or 32 kHz; **Figure 3 and 4**). All ABR and EFR measures were correlated with DPOAE_40_ and DPOAE threshold at both 16 and 32 kHz (**Table 2**). In contrast, evoked measures were only clearly correlated with DPOAE_55_ at 32 kHz, not at 16 kHz (**Table 2**). This may be because high level DP-grams are dominated by passive amplification at 16 kHz, whereas some active amplification from OHCs remains present at 32 kHz (Avan et al., 2003; Mills, 1997).

**Figure 3:**
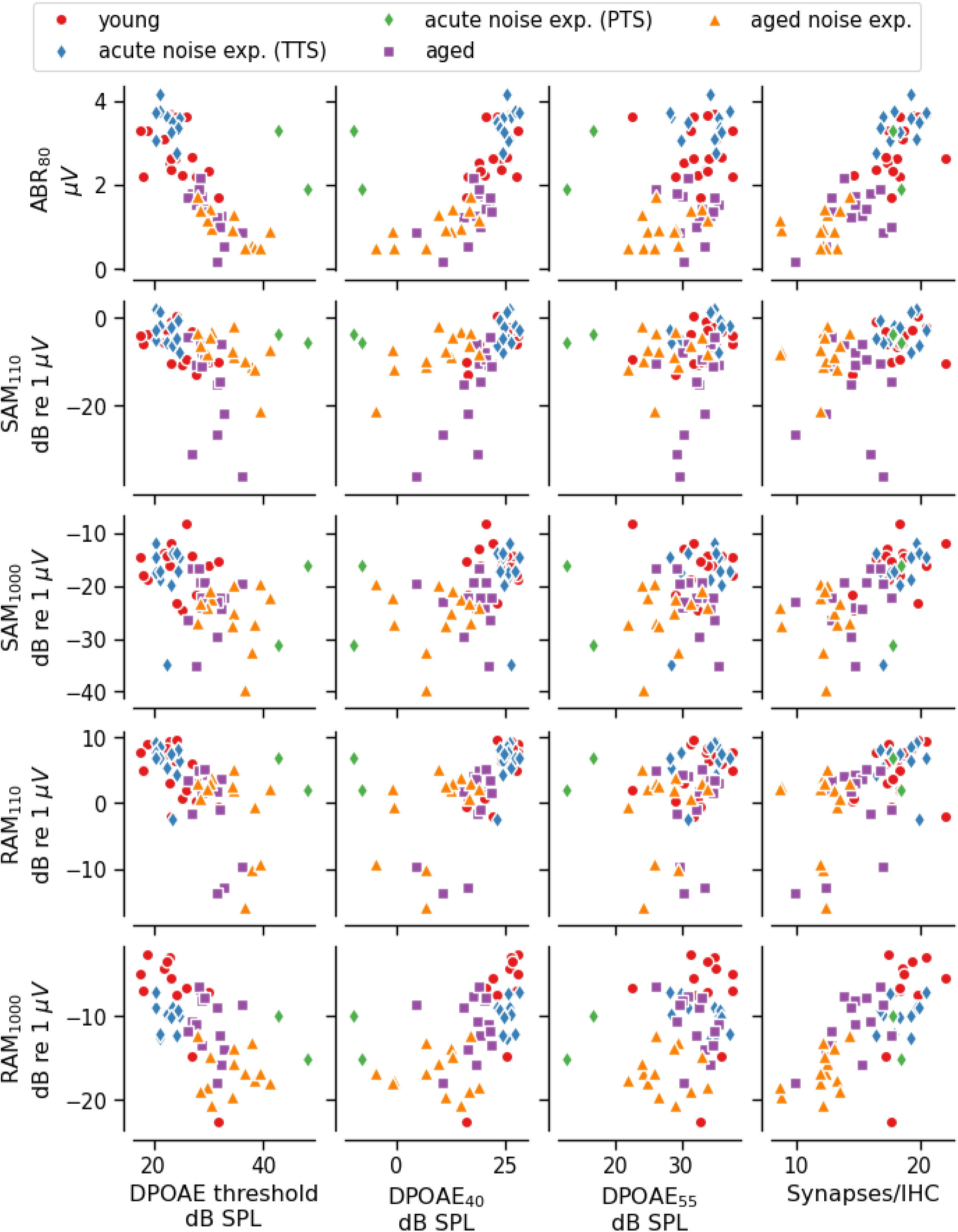
Correlations between evoked potentials and DPOAE measures or synapse counts at 16 kHz. The correlation between each of the evoked potential measures and three DPOAE measures (DPOAE threshold, DPOAE level at *L_2_* = 40 dB SPL [DPOAE_40_], DPOAE level at *L_2_* = 55 dB SPL [DPOAE_55_]) measured for a tone with *f_2_* = 16 kHz or number of synapses per IHC at the 16 kHz tonotopic region. For all plots, symbols indicate data from individual animals and the color/symbol type indicates the experimental group. See Table 2 for correlation coefficients. ABR_80_ = ABR wave 1 amplitude at 80 dB SPL, SAM_110_ = SAM EFR modulated at 110 Hz, SAM_1000_ = SAM EFR modulated at 1000 Hz, RAM_110_ = RAM EFR modulated at 110 Hz, RAM_1000_ = RAM EFR modulated at 1000 Hz.

**Figure 4:**
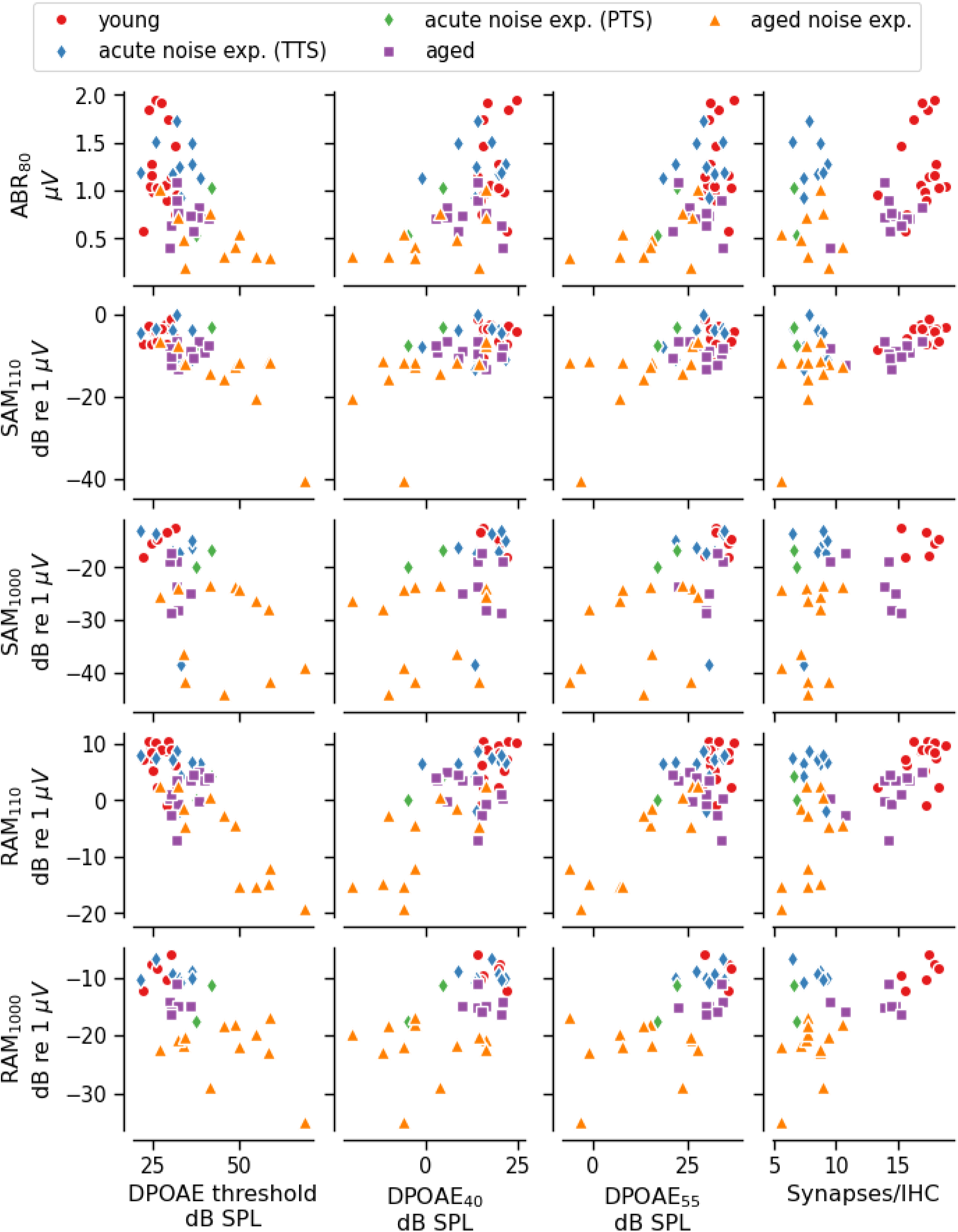
Correlations between evoked potentials and DPOAE measures or synapse counts at 32 kHz. The correlation between each of the evoked potential measures and three DPOAE measures (DPOAE threshold, DPOAE level at *L_2_* = 40 dB SPL, DPOAE level at *L_2_* = 55 dB SPL) measured for a tone with *f_2_* = 32 kHz or number of synapses per IHC at the 32 kHz tonotopic region. For all plots, symbols indicate data from individual animals and the color/symbol type indicates the experimental group. See Table 2 for correlation coefficients.

**Table 2.** Pearson’s correlations between evoked potentials and DPOAE or synapse metrics. The 95% confidence interval is shown in parenthesis. Correlation coefficients are computed using all mice in the study (top rows) or excluding the acute noise exposed group (bottom rows). ABR_80_ = ABR wave 1 amplitude at 80 dB SPL, SAM_110_ = SAM EFR modulated at 110 Hz, SAM_1000_ = SAM EFR modulated at 1000 Hz, RAM_110_ = RAM EFR modulated at 110 Hz, RAM_1000_ = RAM EFR modulated at 1000 Hz, DPOAE_40_ = DPOAE level for *L_2_* = 40 dB SPL, DPOAE_55_ = DPOAE level for *L_2_* = 55 dB SPL.

| Measure | Frequency (kHz) | DPOAE threshold | DPOAE<sub>40</sub> | DPOAE<sub>55</sub> | Synapses/IHC |
| --- | --- | --- | --- | --- | --- |
| All mice |  |  |  |  |  |
| ABR<sub>80</sub> | 16 kHz | -0.7 | 0.61 | 0.26 | 0.8 |
|  |  | (-0.81, -0.53) | (0.42, 0.75) | (0.00, 0.49) | (0.69, 0.88) |
|  | 32 kHz | -0.56 | 0.54 | 0.54 | 0.3 |
|  |  | (-0.73, -0.33) | (0.30, 0.71) | (0.30, 0.71) | (0.01, 0.54) |
| SAM<sub>110</sub> | 16 kHz | -0.38 | 0.37 | 0.14 | 0.31 |
|  |  | (-0.58, -0.13) | (0.13, 0.58) | (-0.12, 0.39) | (0.06, 0.53) |
|  | 32 kHz | -0.72 | 0.55 | 0.62 | 0.42 |
|  |  | (-0.83, -0.55) | (0.32, 0.72) | (0.41, 0.77) | (0.15, 0.62) |
| SAM<sub>1000</sub> | 16 kHz | -0.45 | 0.4 | 0.25 | 0.57 |
|  |  | (-0.64, -0.21) | (0.15, 0.60) | (-0.01, 0.48) | (0.37, 0.73) |
|  | 32 kHz | -0.55 | 0.47 | 0.6 | 0.38 |
|  |  | (-0.75, -0.25) | (0.16, 0.70) | (0.33, 0.78) | (0.05, 0.64) |
| RAM<sub>110</sub> | 16 kHz | -0.54 | 0.52 | 0.27 | 0.49 |
|  |  | (-0.70, -0.32) | (0.30, 0.69) | (0.01, 0.49) | (0.26, 0.67) |
|  | 32 kHz | -0.81 | 0.71 | 0.77 | 0.5 |
|  |  | (-0.89, -0.68) | (0.54, 0.82) | (0.63, 0.86) | (0.27, 0.68) |
| RAM<sub>1000</sub> | 16 kHz | -0.6 | 0.53 | 0.26 | 0.69 |
|  |  | (-0.75, -0.39) | (0.30, 0.70) | (-0.02, 0.50) | (0.51, 0.81) |
|  | 32 kHz | -0.66 | 0.58 | 0.67 | 0.48 |
|  |  | (-0.81, -0.41) | (0.29, 0.76) | (0.42, 0.82) | (0.17, 0.71) |
| Excluding acute noise-exposed |  |  |  |  |  |
| ABR<sub>80</sub> | 16 kHz | -0.83 | 0.77 | 0.37 | 0.79 |
|  |  | (-0.90, -0.71) | (0.61, 0.87) | (0.08, 0.60) | (0.64, 0.88) |
|  | 32 kHz | -0.61 | 0.57 | 0.56 | 0.64 |
|  |  | (-0.78, -0.36) | (0.30, 0.75) | (0.29, 0.75) | (0.40, 0.80) |
| SAM<sub>110</sub> | 16 kHz | -0.4 | 0.44 | 0.18 | 0.19 |
|  |  | (-0.62, -0.12) | (0.16, 0.65) | (-0.12, 0.45) | (-0.12, 0.46) |
|  | 32 kHz | -0.77 | 0.63 | 0.66 | 0.67 |
|  |  | (-0.87, -0.60) | (0.39, 0.79) | (0.43, 0.81) | (0.45, 0.81) |
| SAM<sub>1000</sub> | 16 kHz | -0.52 | 0.43 | 0.2 | 0.62 |
|  |  | (-0.71, -0.26) | (0.15, 0.64) | (-0.10, 0.47) | (0.39, 0.77) |
|  | 32 kHz | -0.58 | 0.51 | 0.67 | 0.66 |
|  |  | (-0.79, -0.23) | (0.14, 0.75) | (0.38, 0.84) | (0.36, 0.84) |
| RAM<sub>110</sub> | 16 kHz | -0.61 | 0.64 | 0.35 | 0.44 |
|  |  | (-0.77, -0.39) | (0.42, 0.79) | (0.06, 0.58) | (0.17, 0.65) |
|  | 32 kHz | -0.84 | 0.76 | 0.81 | 0.79 |
|  |  | (-0.91, -0.72) | (0.59, 0.87) | (0.67, 0.90) | (0.64, 0.89) |
| RAM<sub>1000</sub> | 16 kHz | -0.69 | 0.62 | 0.33 | 0.74 |
|  |  | (-0.83, -0.48) | (0.38, 0.78) | (0.01, 0.58) | (0.55, 0.85) |
|  | 32 kHz | -0.67 | 0.55 | 0.65 | 0.84 |
|  |  | (-0.84, -0.37) | (0.20, 0.78) | (0.35, 0.83) | (0.66, 0.93) |

### Relationship between synapse numbers and ABR/EFR measurements

All evoked potential measures were correlated with synapse numbers at 16 kHz (**Figure 3**, **Table 2**) and 32 kHz (**Figure 4**, **Table 2**). To assess how inclusion of the acute noise exposed mice, who had a large frequency-specific reduction in synapse numbers at 32 kHz, but near-normal evoked potential magnitudes, affected the results, we recomputed the correlations without the acute noise exposed group (**Table 2**).

Recomputing the correlations without the acute noise exposed group resulted in stronger correlations between synapse numbers and all five evoked potential measures at 32 kHz. Notably, excluding the acute noise exposed group resulted in poorer correlations between synapse numbers and two evoked potential measures, SAM_110_ and RAM_110_, but only at 16 kHz. When considering the subset of data representing animals without acute noise exposure, ABR_80_ had the strongest correlation with synapse numbers at 16 kHz and RAM_1000_ had the strongest correlations with synapse number at 32 kHz (**Table 2**), suggesting that ABR_80_ and RAM_1000_ may be more robust predictors of synapse number than the other evoked measures.

### Analysis of the ability of each evoked potential measure to predict synapse numbers

To assess which evoked potential measures have the best ability to predict cochlear synapse numbers, we used linear regression models to predict synapse numbers at cochlear frequencies of 16 and 32 kHz. Since evoked potentials can be affected by OHC dysfunction, the linear regression models optionally included an adjustment for one of the DPOAE measures. Due to the collinearity between evoked potentials and DPOAEs, the actual values of the regression coefficients (i.e., 𝛽 in the equations) cannot be interpreted when more than one predictor is included in the model. However, the predictive power of these models is not affected by multicollinearity (Kutner et al., 2005). Repeated k-fold cross validation was used to estimate the prediction error of each model, allowing for direct comparison of the performance of different combinations of predictors. Data from both frequencies (16 kHz and 32 kHz) was pooled for the model predictions (i.e., frequency-specific effects were not included).

All models overestimated the number of synapses at 32 kHz for the mice with focal synaptopathy due to acute noise exposure (open green circles in **Figure 5**).

**Figure 5:**
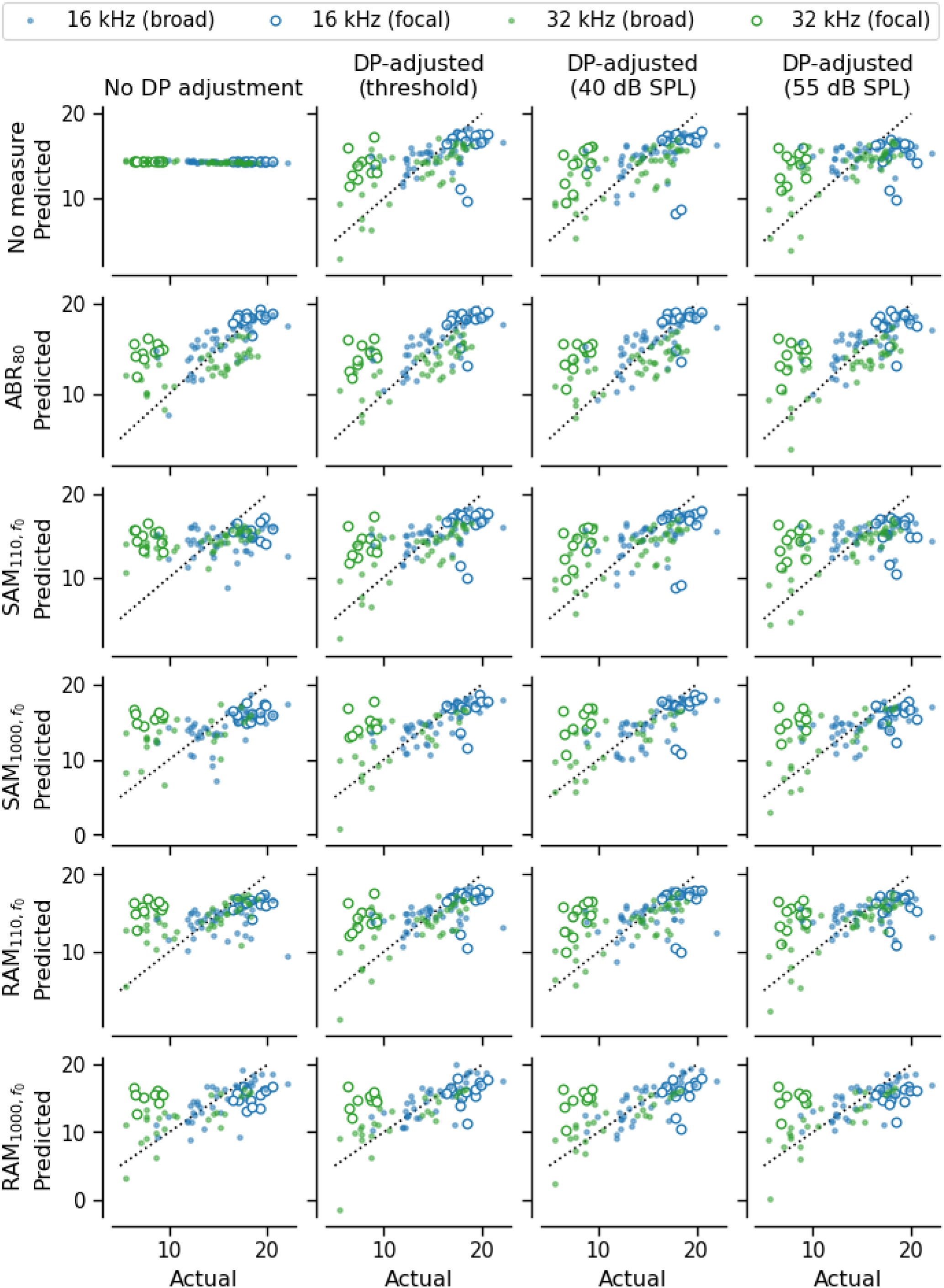
Synapse prediction model accuracy. For each model, the predicted vs. actual number of synapses per IHC is shown. Color indicates the frequency, open circles indicate data from ears with acute noise exposure (focal synaptopathy), and filled circles indicate data from all other ears (no synaptopathy or broad synaptopathy). Dotted line indicates unity. Each row shows the results of the indicated predictor (e.g., ABR wave 1 amplitude at 80 dB SPL [ABR_80_]) without DPOAE adjustment (first column) or adjusted using one of three DPOAE measures (columns 2-4). The first plot in the first row shows the results of an intercept-only model, which was used to establish baseline performance. In this model, the predicted synapse count is the average synapse count for the whole sample. The remaining subplots in the first row show the results of predicting synapses using only DPOAE metrics. These models establish a reference against which the ABR and EFR predictors can be evaluated. SAM and RAM EFR magnitude were calculated as the EFR power at the modulation frequency without correction for the noise floor (i.e., f_0_, see Figure 1). See Table 3 for correlation coefficients.

**Table 3.** Pearson’s correlations between predicted and actual synapse numbers for each linear regression model. The 95% confidence interval is shown in parenthesis. Correlation coefficients are computed using all mice in the study (top rows) or excluding the acute noise exposed group (bottom rows). Columns show the DPOAE adjustment that was used.

| Measure | No adjustment |  | DPOAE threshold |  | DPOAE<sub>40</sub> |  | DPOAE<sub>55</sub> |  |
| --- | --- | --- | --- | --- | --- | --- | --- | --- |
| All mice |  |  |  |  |  |  |  |  |
| None |  |  | 0.49 | (0.47, 0.52) | 0.44 | (0.41, 0.47) | 0.34 | (0.31, 0.37) |
| ABR<sub>80</sub> | 0.73 | (0.72, 0.75) | 0.69 | (0.67, 0.71) | 0.69 | (0.67, 0.71) | 0.66 | (0.64, 0.68) |
| SAM<sub>110</sub> | 0.23 | (0.20, 0.26) | 0.49 | (0.47, 0.52) | 0.45 | (0.42, 0.47) | 0.36 | (0.33, 0.39) |
| SAM<sub>1000</sub> | 0.28 | (0.24, 0.31) | 0.51 | (0.48, 0.54) | 0.47 | (0.44, 0.50) | 0.38 | (0.35, 0.41) |
| RAM<sub>110</sub> | 0.28 | (0.25, 0.31) | 0.48 | (0.45, 0.50) | 0.44 | (0.41, 0.46) | 0.36 | (0.33, 0.39) |
| RAM<sub>1000</sub> | 0.26 | (0.23, 0.30) | 0.45 | (0.42, 0.48) | 0.42 | (0.39, 0.45) | 0.34 | (0.31, 0.37) |
| Excluding acute noise-exposed |  |  |  |  |  |  |  |  |
| None |  |  | 0.68 | (0.64, 0.71) | 0.64 | (0.60, 0.68) | 0.63 | (0.59, 0.67) |
| ABR<sub>80</sub> | 0.68 | (0.65, 0.72) | 0.72 | (0.68, 0.75) | 0.7 | (0.67, 0.74) | 0.73 | (0.70, 0.77) |
| SAM<sub>110</sub> | 0.29 | (0.22, 0.35) | 0.66 | (0.62, 0.70) | 0.63 | (0.59, 0.67) | 0.63 | (0.58, 0.67) |
| SAM<sub>1000</sub> | 0.57 | (0.52, 0.62) | 0.77 | (0.73, 0.79) | 0.78 | (0.75, 0.81) | 0.74 | (0.70, 0.77) |
| RAM<sub>110</sub> | 0.5 | (0.45, 0.55) | 0.65 | (0.61, 0.69) | 0.64 | (0.59, 0.67) | 0.64 | (0.60, 0.68) |
| RAM<sub>1000</sub> | 0.78 | (0.74, 0.80) | 0.82 | (0.80, 0.85) | 0.84 | (0.82, 0.86) | 0.82 | (0.79, 0.84) |

However, the models were generally successful at predicting synapse numbers for mice with no synaptopathy or broad synaptopathy regardless of frequency (dots in **Figure 5**). When excluding the acute noise exposed group, the RAM_1000_ showed the strongest correlation between predicted and actual synapse counts as compared to the other measures regardless of statistical adjustment for DPOAEs (bottom row of **Figure 5**, **Table 3**). When considering the full dataset (i.e., including the acute noise exposed group), the ABR_80_ showed the strongest correlation between predicted and actual synapse count regardless of statistical adjustment for DPOAEs (second row of **Figure 5**, **Table 3**). This suggests that, while RAM_1000_ is the strongest predictor of broad synaptopathy, ABR_80_ is the strongest predictor of synapse number when there are cases of focal synaptopathy.

Since correlations simply capture linear relationships between variables, the prediction accuracy of each model was quantified using the root mean squared error (RMSE). The RMSE values indicate the prediction error in terms of number of synapses, with smaller values indicating better predictive power. RMSE was computed separately for each repeat and fold and then the mean and standard error were computed across all repeats and folds to give the pointwise estimate and standard error of the expected RMSE for future data. Baseline performance was assessed using an intercept-only model in which the predicted synapse count was the average synapse count for the entire dataset (blue bar in **Figure 6A**). All models incorporating DPOAE and/or evoked potential data performed better than baseline when predicting the number of synapses in mice with no synaptopathy or broad synaptopathy (i.e., excluding the acute noise exposed group, **Figure 6A**). In contrast, when predicting the number of synapses in mice with focal synaptopathy, only ABR_80_ clearly performed better than chance regardless of whether DPOAE adjustment was included (**Figure 7**). In the mice with focal synaptopathy, all the EFR measures performed only slightly better than chance when predicting cochlear synapses and only when adjusted for DPOAE threshold or DPOAE_40_. However, the EFR measures, when adjusted for DPOAEs, did not outperform the matched DPOAE-only model. This indicates that the predictions generated by the EFR models are primarily driven by the inclusion of the DPOAE adjustment and that the EFR measurements likely did not contribute to the prediction.

**Figure 6:**
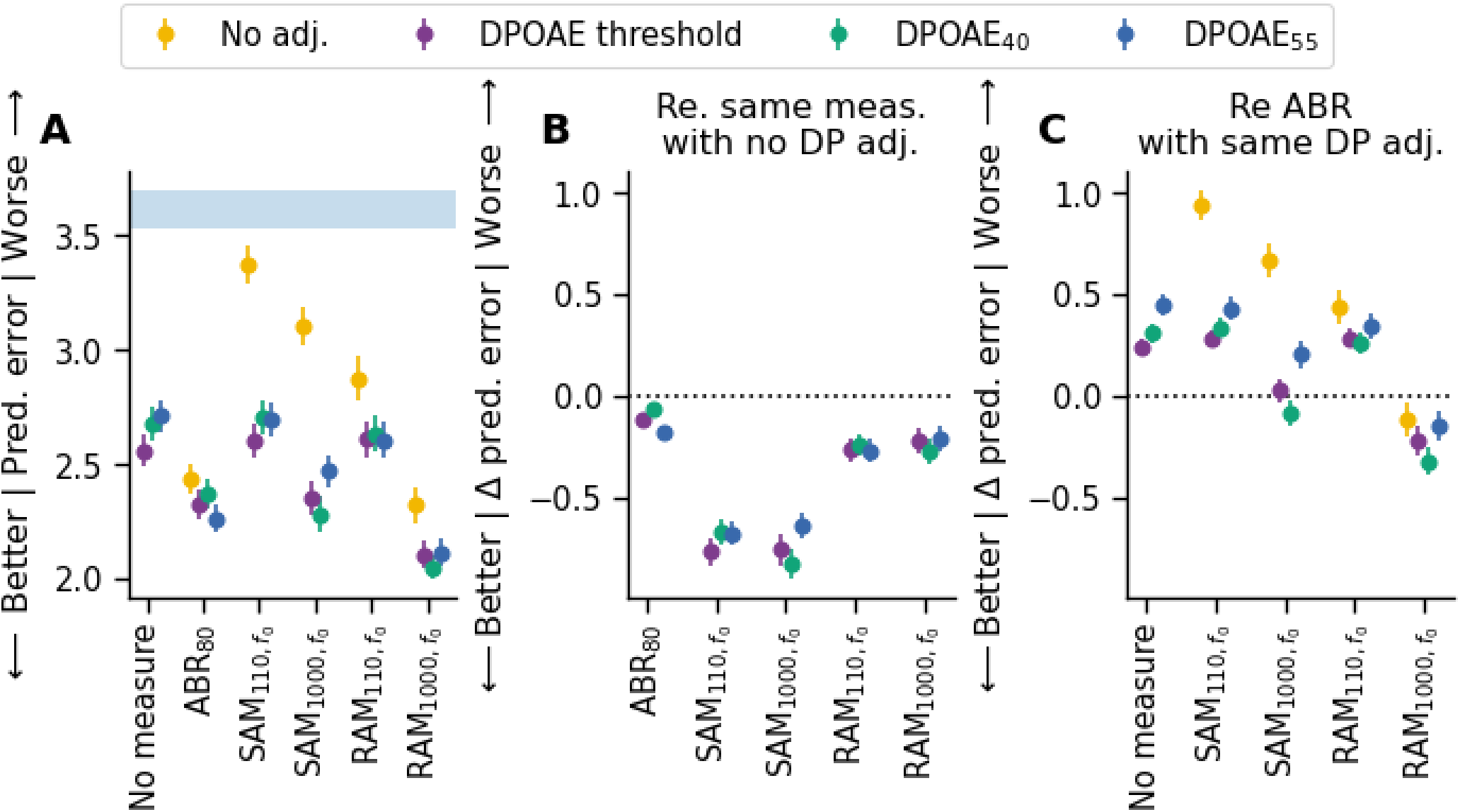
Relative ability of evoked potential models to predict synapse numbers in mice with no synaptopathy or broad synaptopathy. **(A)** Root mean square error (RMSE) of each evoked potential model with and without DPOAE adjustment. Shaded blue region indicates baseline performance +/- standard error of the mean (SEM) for a model with no predictors (intercept-only model where the predicted value was set to the mean synapse number for the sample). **(B)** Change in prediction error for each model, after DPOAE adjustment, relative to the same model without DPOAE adjustment. **(C)** Change in prediction error for each model relative to the ABR_80_ model when matched for DPOAE adjustment (i.e., RAM_1000_ model adjusted for DPOAE threshold would be compared to ABR_80_ model adjusted for DPOAE threshold). For all panels, the RMSE was computed on the data pooled across 16 and 32 kHz only for mice in the young, aged after noise exposure, and aged groups. Markers indicate the RMSE of the corresponding model averaged across all repeats and folds, colors indicate the DPOAE adjustment applied, and error bars indicate the SEM of the RMSE across all repeats and folds. SAM and RAM EFR magnitude were calculated as the EFR power at the modulation frequency without correction for the noise floor (i.e., f_0_, see Figure 1).

**Figure 7:**
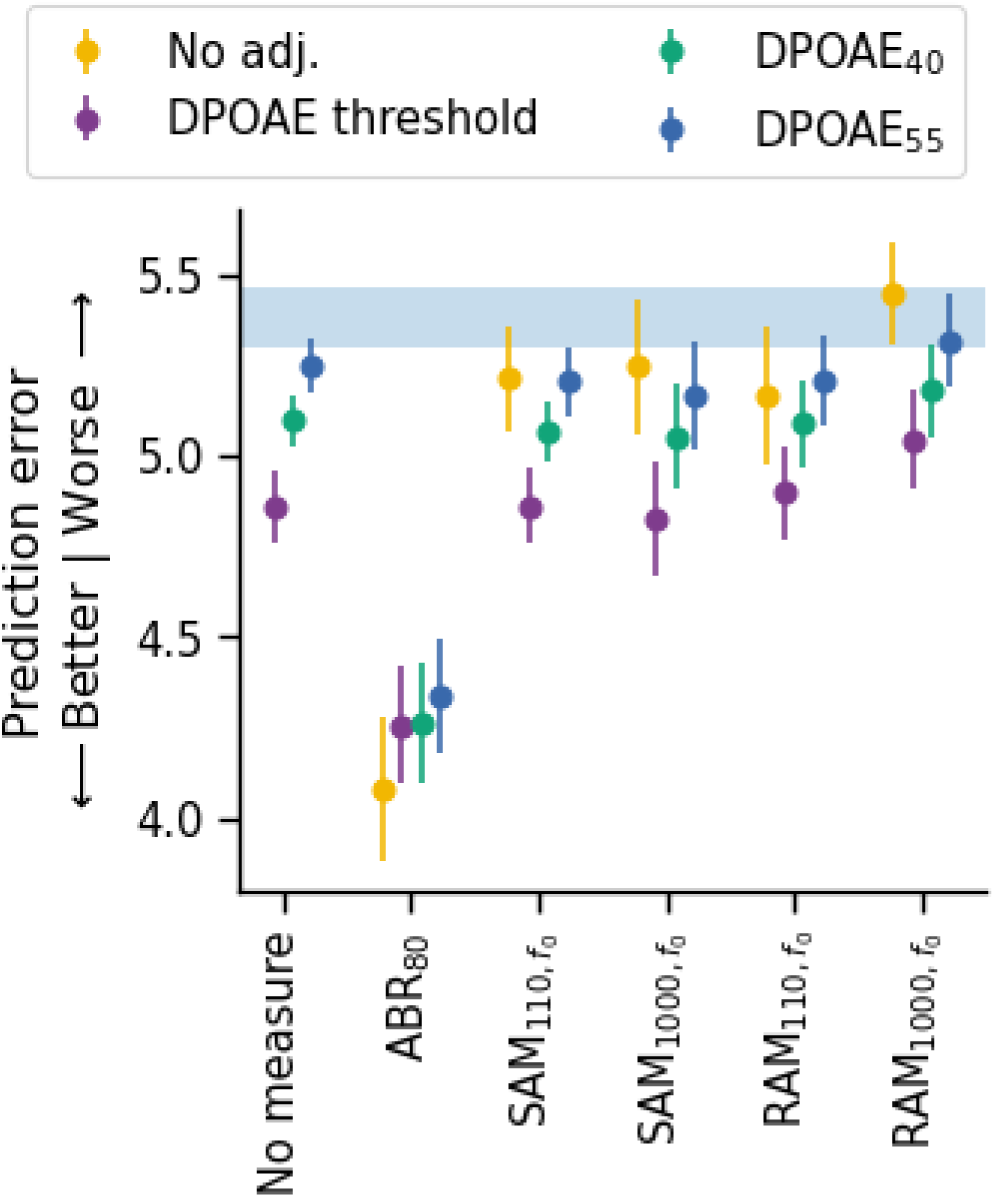
Relative ability of evoked potential models to predict synapse numbers in mice with focal synaptopathy. Root mean square error (RMSE) of each evoked potential model with and without DPOAE adjustment. Shaded blue region indicates baseline performance +/- standard error of the mean (SEM) of a model with no predictors (intercept-only model where the predicted value was set to the mean synapse number for the sample). The RMSE was computed on the data pooled across 16 and 32 kHz only for mice in the acute noise exposed group. Markers indicate the RMSE of the corresponding model averaged across all repeats and folds, colors indicate the DPOAE adjustment applied, and error bars indicate the SEM of the RMSE across all repeats and folds. SAM and RAM EFR magnitude were calculated as the EFR power at the modulation frequency without correction for the noise floor (i.e., f_0_, see Figure 1).

As an additional test of model performance, we fit each model to the full dataset without cross-validation and calculated the corrected Akaike Information Criterion (AICc). Comparisons were made across the full set of models (**Table 4**). These results demonstrate that ABR_80_ corrected for DPOAE_55_ is likely the best predictor of synapse count when considering both mice with and without focal synaptopathy. However, the set of ABR_80_ models that used other forms of DPOAE adjustment or did not adjust for DPOAEs performed comparably well as indicated by ΔAICc <= 2.8. We recomputed the AICc after fitting each model with the acute noise exposed group excluded (**Table 4, bottom rows**). This analysis corroborated the predictions of our cross-validation that RAM_1000_ adjusted for DPOAE threshold significantly outperformed all other evoked measures; however, the RAM_1000_ models incorporating other forms of DPOAE adjustment should not be completely rejected since they had ΔAICc <= 6.37. Given the poor performance of the evoked potentials in predicting focal synaptopathy, all subsequent analyses excluded the mice from the acute noise exposed group. Thus, the following results are specific to cases where no synaptopathy or broad synaptopathy would be expected.

**Table 4.** ΔAICc values for each model compared to the best performing overall model. ΔAICc is computed using all mice in the study (top rows) or excluding the acute noise exposed group (bottom rows). The best performing model in each case is indicated by a dash. Models with ΔAICc values ≤ 2 generally are considered comparable to the best performing model whereas models with ΔAICc values > 10 are considered to have poorer performance than the best performing model. Models with ΔAICc between 2 and 10 should not be rejected outright even though the evidence suggests that the model with a lower AICc is better.

| DP adjustment | None | ABR<sub>80</sub> | SAM<sub>110</sub> | SAM<sub>1000</sub> | RAM<sub>110</sub> | RAM<sub>1000</sub> |
| --- | --- | --- | --- | --- | --- | --- |
| All mice |  |  |  |  |  |  |
| None | 59.44 | 2.8 | 56.77 | 44.14 | 50.96 | 28.17 |
| DPOAE threshold | 24.48 | 0.22 | 26.58 | 16.73 | 26.57 | 7.26 |
| DPOAE<sub>40</sub> | 31.69 | 2.33 | 33.71 | 21.35 | 33.6 | 13.48 |
| DPOAE<sub>55</sub> | 38.19 | - | 39.15 | 27.49 | 37.9 | 16.73 |
| Excluding acute noise-exposed |  |  |  |  |  |  |
| None | 80.9 | 34.27 | 80.38 | 58.67 | 72.64 | 27.76 |
| DPOAE threshold | 42.41 | 27.82 | 44.17 | 25.89 | 44.53 | - |
| DPOAE<sub>40</sub> | 47.33 | 30.66 | 48.84 | 28.69 | 49.46 | 6.37 |
| DPOAE<sub>55</sub> | 52.1 | 25.37 | 54.09 | 34.01 | 52.24 | 5.67 |

The robustness of each evoked potential measure in the presence of OHC dysfunction varied, as indicated by the difference in RMSE between models with no DPOAE adjustment versus models incorporating a DPOAE adjustment. To quantify the effect of adjusting for DPOAEs, the difference in prediction error for each evoked potential model with DPOAE adjustment was compared against the same evoked potential model without DPOAE adjustment (**Figure 6B**). DPOAE adjustment improved the predictive performance of models for each evoked potential measure, with the greatest impact on the SAM_110_ (0.66 to 0.77 improvement) and SAM_1000_ (0.64 to 0.84 improvement) models. The impact of DPOAE adjustment was intermediate for the RAM_110_ (0.24 to 0.29 improvement) and RAM_1000_ (0.21 to 0.27 improvement) models. DPOAE adjustment only had marginal impact for ABR_80_ (0.07 to 0.17 improvement).

As an additional test of the value of the DPOAE adjustment, we compared the AICc across the prediction models for a given evoked potential measure based on the DPOAE adjustment that was used (**Table 5**). Except for ABR_80,_ all other measures performed better at predicting synapse numbers when adjusted for DPOAE threshold. Although ABR_80_ performed best when adjusted for DPOAE_55_, ΔAICc was <= 8.9 regardless of if, or how, ABR_80_ was adjusted for DPOAEs. Notably, all other evoked measures had much better AICc scores when adjusted for DPOAEs as compared to no adjustment, providing additional evidence that the ABR is a relatively robust predictor of synapse numbers in the presence of OHC dysfunction. In the DPOAE-only, SAM_110_, SAM_1000_, RAM_110_, and RAM_1000_ models, there was a clear trend of model performance decreasing with increasing DPOAE level, although ΔAICc was <= 9.69 in all cases. This indicates that assessing DPOAEs using lower *L_2_* yields better results, possibly because high level DP-grams are more likely to reflect passive amplification (Avan et al., 2003; Mills, 1997).

**Table 5.** ΔAICc values for each method of adjusting for DPOAEs relative to the best performing model for each evoked potential. ΔAICc is computed across the models for a given evoked potential to assess which DPOAE adjustment method performs best for a particular evoked potential measure (i.e., comparisons should be made across columns, not across rows). The best performing DPOAE adjustment for each evoked potential model is indicated by a dash.

| Measure | No adjustment | DPOAE<br>threshold | DPOAE<sub>40</sub> | DPOAE<sub>55</sub> |
| --- | --- | --- | --- | --- |
| None | 38.48 | - | 4.91 | 9.69 |
| ABR<sub>80</sub> | 8.9 | 2.44 | 5.29 | - |
| SAM<sub>110</sub> | 36.21 | - | 4.67 | 9.92 |
| SAM<sub>1000</sub> | 32.78 | - | 2.81 | 8.12 |
| RAM<sub>110</sub> | 28.12 | - | 4.94 | 7.71 |
| RAM<sub>1000</sub> | 27.76 | - | 6.37 | 5.67 |

Given that ABR wave 1 amplitude is the gold standard for assessing auditory nerve function, the predictive performance of each model was compared with the ABR wave 1 amplitude model (**Figure 6C**). For all comparisons, the DP adjustment was matched with that used in the ABR model (i.e., the SAM_110_ model adjusted for DPOAE threshold was compared with the ABR_80_ model adjusted for DPOAE threshold).

Compared to ABR_80_, the prediction error was worse for the DPOAE-only (0.24 to 1.18 worse), SAM_110_ (0.28 to 0.94 worse), and RAM_110_ (0.27 to 0.44 worse) models, regardless of which method of DPOAE adjustment was used. In contrast, the SAM_1000_ model performed similarly to the ABR_80_ model (−0.09 to 0.67 worse), but only when adjusted for DPOAE_40_ or DPOAE threshold. The RAM_1000_ model outperformed the ABR_80_ (0.12 to 0.32 better), particularly when adjusted for DPOAE_40_.

As an additional test of the performance of the RAM_1000_ model, we compared each of the models using AICc (**Table 6**). Comparisons were made across all evoked potential models for a given DPOAE adjustment. Regardless of the DPOAE adjustment used, the RAM_1000_ model significantly outperformed all the other models except for the ABR_80_ model which performed only slightly worse than the RAM_1000_ model when there was no DPOAE adjustment (ΔAICc = 6.51).

**Table 6.** ΔAICc values for each evoked potential model relative to the best performing evoked potential model for a given DPOAE adjustment. ΔAICc is computed across the evoked potential models for a given DPOAE adjustment method to assess which evoked potential model is the best predictor of synapse count (i.e., comparisons should be made across columns, not across rows). The best performing evoked potential for each DPOAE adjustment is indicated by a dash.

| DP adjustment | None | ABR<sub>80</sub> | SAM<sub>110</sub> | SAM<sub>1000</sub> | RAM<sub>110</sub> | RAM<sub>1000</sub> |
| --- | --- | --- | --- | --- | --- | --- |
| No adjustment | 53.14 | 6.51 | 52.62 | 30.91 | 44.88 | - |
| DPOAE threshold | 42.41 | 27.82 | 44.17 | 25.89 | 44.53 | - |
| DPOAE<sub>40</sub> | 40.95 | 24.29 | 42.46 | 22.32 | 43.09 | - |
| DPOAE<sub>55</sub> | 46.44 | 19.71 | 48.42 | 28.34 | 46.57 | - |

### Effect of EFR processing on synapse prediction error

The impact of different methods of calculating EFR magnitude (i.e., calculating EFR power relative to the noise floor [NF], including harmonics in the sum [f_0-4_], or using phase-locking value [PLV]) on the ability of the EFR models to predict synapse numbers was evaluated (**Figure 8**). For this comparison, all EFR models were compared to the prediction error for the absolute EFR magnitude at the modulation frequency (i.e., f_0_).

**Figure 8:**
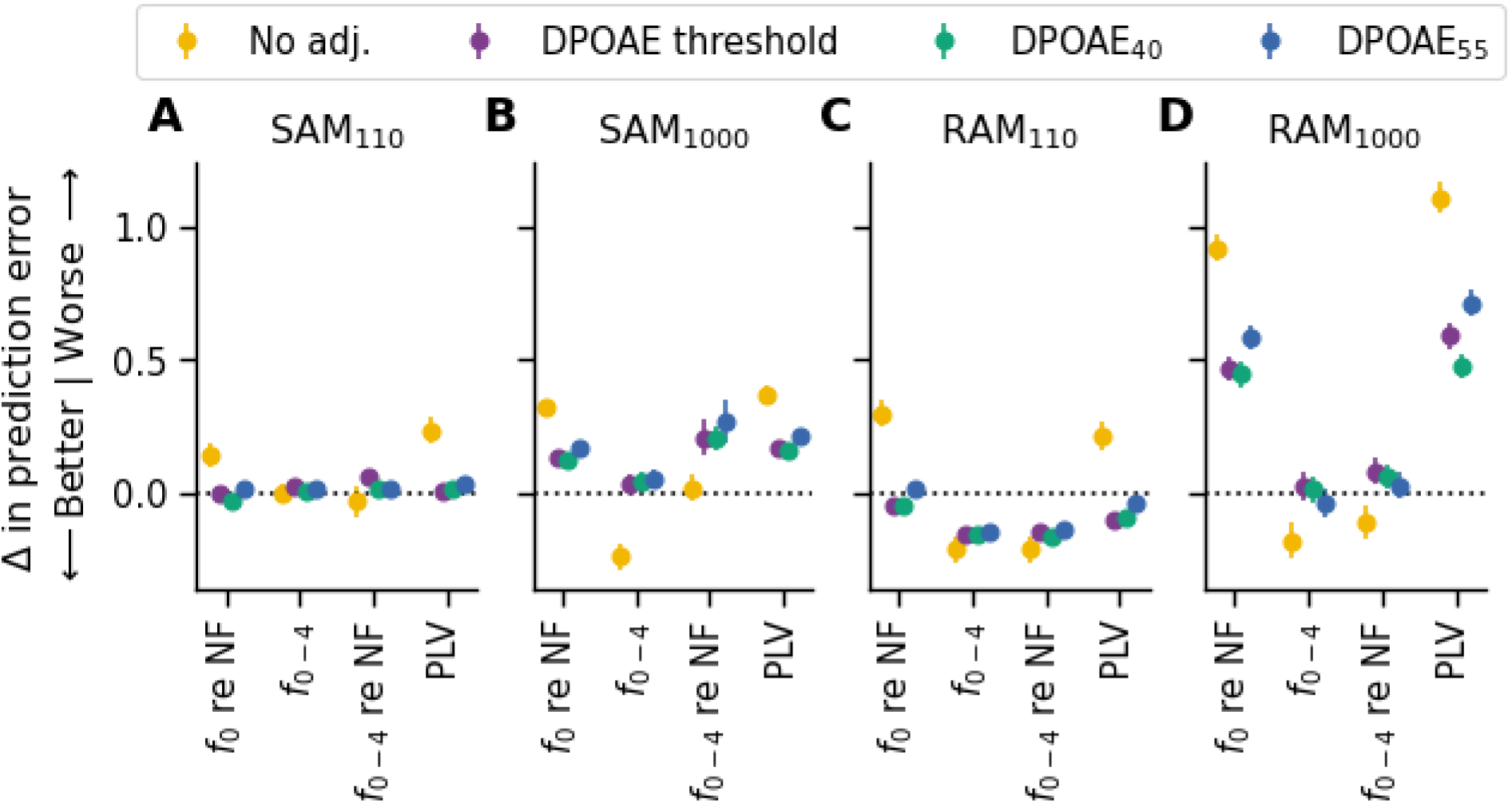
Impact of EFR processing on synapse prediction performance in mice with no synaptopathy or broad synaptopathy. EFR model results are plotted relative to the model where the same EFR measure was calculated using only the power at the modulation frequency without correction for the noise floor (f_0_, see Figure 1). The change in root mean square error (RMSE) is plotted for the **(A)** SAM_110_, **(B)** SAM_1000_, **(C)** RAM_110_, and **(D)** RAM_1000_ models. The RMSE was computed on the data pooled across 16 and 32 kHz only for mice in the young, aged noise exposed, and aged groups. Markers indicate the change in RMSE of the corresponding model averaged across all repeats and folds, colors indicate the DPOAE adjustment applied, and error bars indicate the standard error of the mean (SEM) of the change in RMSE across all repeats and folds. Dashed line indicates the reference model (f_0_). PLV = phase locking value, NF = noise floor.

For the SAM_110_ model, there was very little difference between the different methods of calculating EFR magnitude (**Figure 8A**). In contrast, for the SAM_1000_ model, referencing the EFR magnitude to the noise floor (either f_0_ re NF or f_0-4_ re NF) or using the PLV resulted in slightly decreased model performance compared to using f_0_ (**Figure 8B**). For RAM_110_, the models that used f_0_-f_4_, with or without incorporating the NF (f_0-4_ and f_0-4_ re NF) performed better than the reference model using f_0_ (**Figure 8C**). For RAM_1000_, the f_0-4_ models (with or without incorporating NF) performed similarly to the reference model (f_0_), while the f_0_ re NF and PLV models performed more poorly (**Figure 8D**). Across all EFR stimuli, summing the EFR power at multiples of the modulation frequency (f_0-4_) resulted in similar or better performance than using only the power at the modulation frequency (f_0_), indicating that there is no disadvantage to summing EFR power at multiples of the modulation frequency. When summing EFR power at multiples of the modulation frequency, referencing the sum to the noise floor (i.e., f_0-4_ re NF) resulted in poorer performance only for the SAM_1000_ models. Thus, the most uniformly consistent performing analysis across all EFR stimuli was the sum of the EFR power at multiples of the modulation frequency without referencing to the noise floor (f_0-4_). Since the noise floor reflects other factors (e.g., arousal state, electrical interference, etc.), incorporating the noise floor into the calculations may increase measurement error.

### Effect of ABR stimulus level on synapse prediction error

High intensity level ABR stimuli are less frequency-specific than lower intensity stimuli due to the spread of excitation along the cochlear partition (Kiang, 1965). For this reason, the analysis was repeated using ABR wave 1 amplitude models for lower stimulus levels (40 to 70 dB SPL; [ABR_40_ to ABR_70_]). Regardless of the DPOAE adjustment used, the models using lower stimulus levels resulted in lower prediction errors than the ABR_80_ model (0.13 to 0.46 improvement; **Figure 9A**). However, the lower stimulus levels come with the trade-off of missing data. The number of animals in the prediction model decreased from 44 (for 70- and 80-dB SPL) to 33 at 40 dB SPL. To avoid excluding data points at lower ABR stimulus levels, an alternate metric for assessing wave 1 amplitude was used where the maximum and minimum values between 1 and 2.4 msec (the range of wave 1 latencies observed in this study) were calculated from the ABR waveform (i.e., peak to peak [PTP]). Wave 1 amplitude was calculated in this fashion for all stimulus levels even if they were subthreshold. This reassessment showed that ABR wave 1 amplitudes at levels of 60-70 SPL, but not 40-50 dB SPL, yielded lower prediction errors than 80 dB SPL (**Figure 9B**).

**Figure 9:**
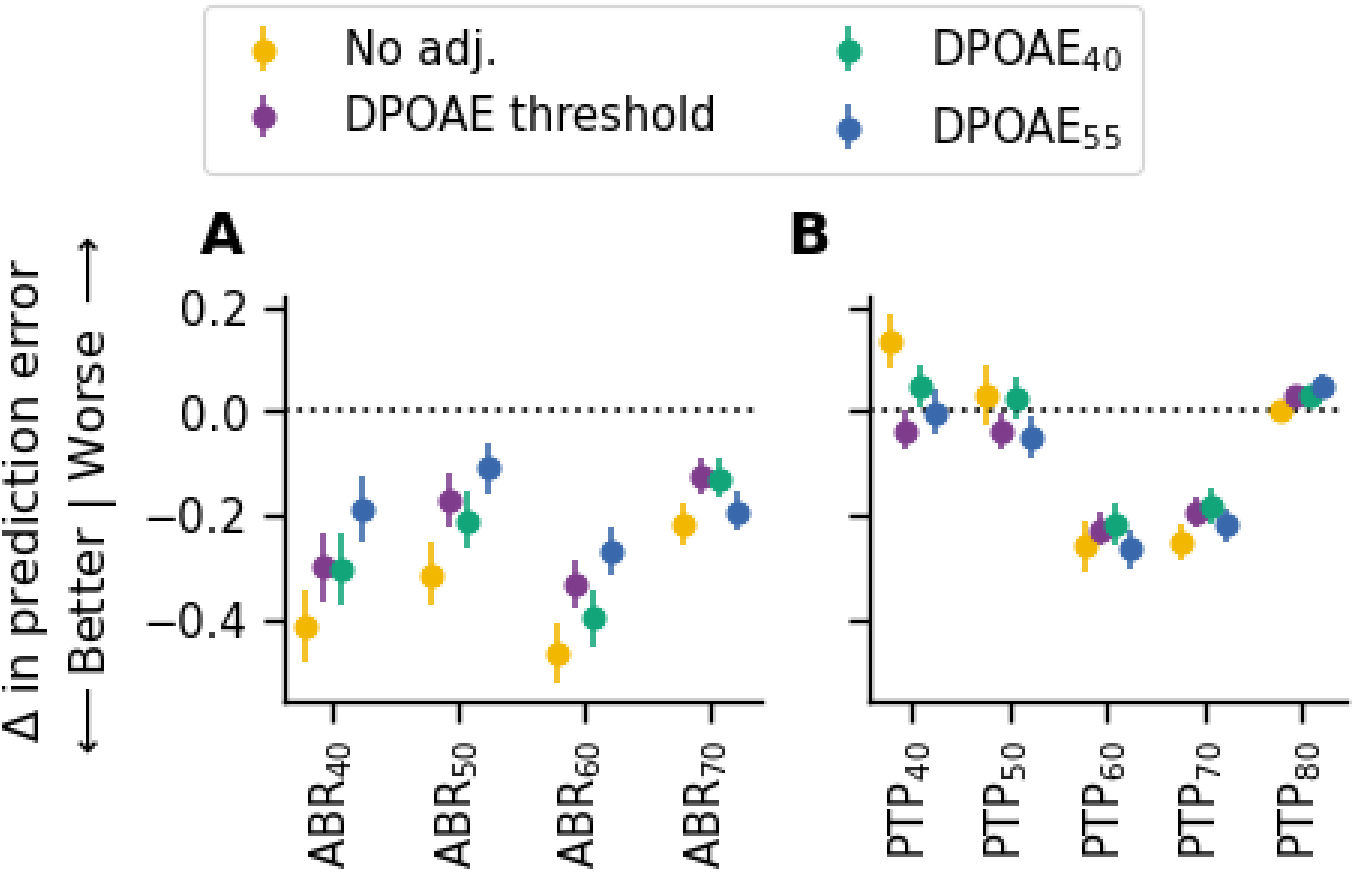
Impact of ABR stimulus level on synapse prediction performance in mice with no synaptopathy or broad synaptopathy. **(A)** Change in prediction error for ABR wave 1 amplitude models for intensity levels of 40-70 dB SPL (ABR_40_, ABR_50_, ABR_60_ and ABR_70_) relative to the ABR wave 1 amplitude model for 80 dB SPL (ABR_80_) when matched for DPOAE adjustment. **(B)** Change in prediction error when calculating ABR wave 1 amplitude using an automated algorithm for each intensity level (PTP_40_, PTP_50_, PTP_60_, PTP_70,_ and PTP_80_) relative to ABR_80_. Wave 1 amplitude was calculated as the difference between the minimum and maximum values (i.e., peak to peak, or PTP) in a window centered around the expected ABR wave 1 latency. For all panels, the change in root mean square error (RMSE) was computed on the data pooled across 16 and 32 kHz only for mice in the young, aged noise exposed, and aged groups. Markers indicate the change in RMSE for the corresponding model relative to ABR_80_ averaged across all repeats and folds, colors indicate the DPOAE adjustment applied, and error bars indicate the standard error of the mean (SEM) of the change in RMSE across all repeats and folds. The dashed line in both panels indicates the reference model (ABR_80_).

### Comparison of models with single evoked potential measures and models with multiple measures

Since each evoked potential may encode different information about auditory nerve function, we tested whether combining multiple evoked potential measures may improve the ability to predict synapse numbers relative to the highest performing single evoked potential model (RAM_1000_). Prior to starting the analysis, we established several pre-determined combinations that we intended to test. Of these pre-determined combinations, a model including both ABR_80_ and RAM_1000_ showed the biggest improvement over the RAM_1000_ model (**Figure 10**). After reviewing the results of our initial analysis, which suggested that the ABR_60_ model outperforms the ABR_80_ model (**Figure 9**) we added a new combination for testing (asterisks in **Figure 10**). This new analysis revealed that the best performance was observed for the model that included both RAM_1000_ and ABR_60_ when no DPOAE adjustment was applied, yielding improvements over the ABR_60_-only (ΔAICc = 21.13 to 23.75) and RAM_1000_-only (ΔAICc = 31.78 to 59.54) models regardless of whether these single-measure models were adjusted for DPOAEs (**Table 7**). However, incorporating a DPOAE adjustment in the combined RAM_1000_ and ABR_60_ model did not significantly alter model performance (ΔAICc between 1.31 and 2.21 relative to the combined model without DPOAE adjustment). **Figure 10** also shows the performance of two alternate single measure models relative to the RAM_1000_ model. The RAM_1000_ model outperformed the RAM_110,_ f_0-4_ model (ΔAICc = 34.71 to 43.89). The ABR_60_ model performed slightly better than the RAM_1000_ model (ΔAICc = 8.03 to 37.95), particularly when no DPOAE adjustment was used (ΔAICc = 37.95), but not better than the combined RAM_1000_ and ABR_60_ model.

**Figure 10:**
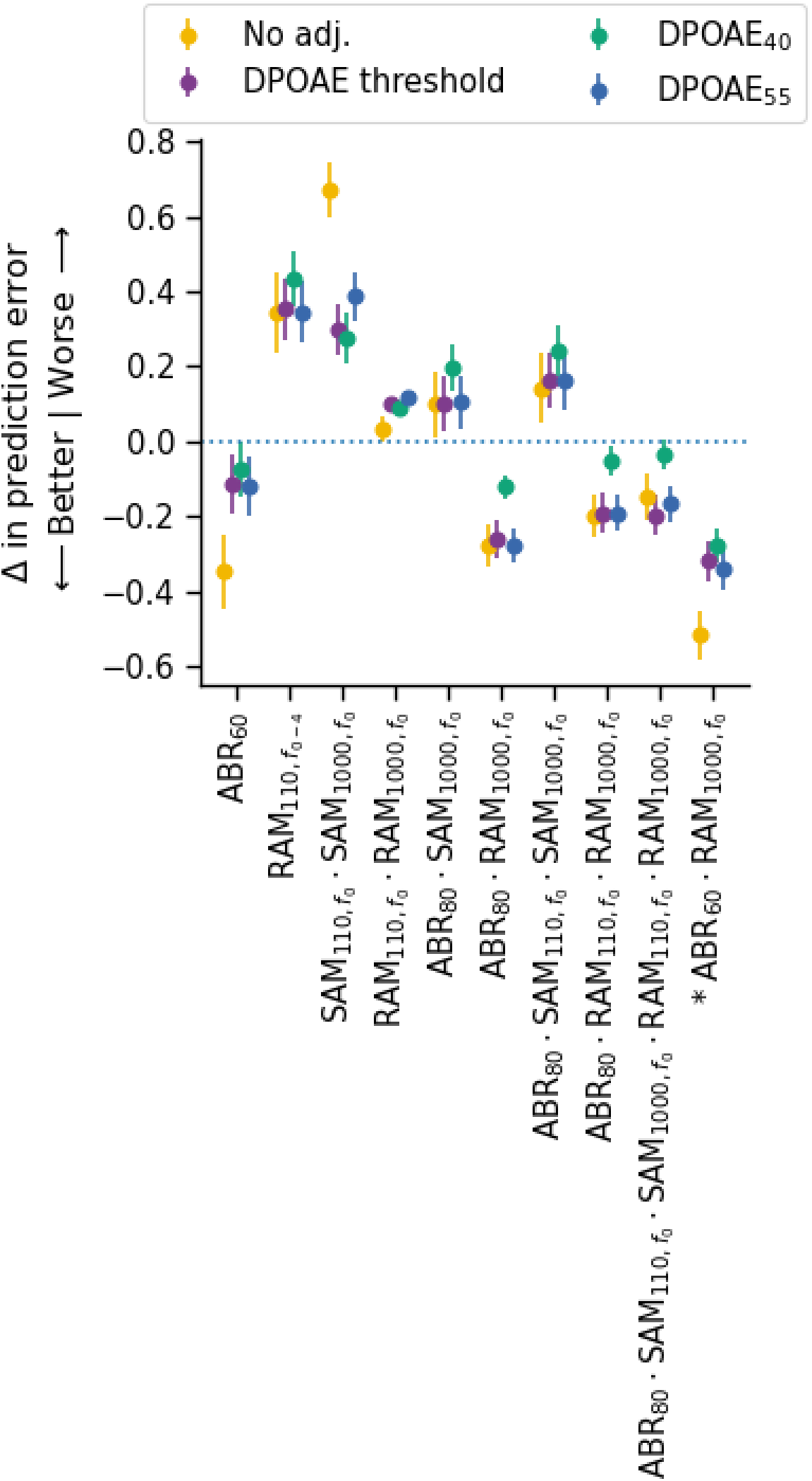
Impact on synapse prediction performance of incorporating multiple evoked potential measures in mice with no synaptopathy or broad synaptopathy. Change in prediction error for models incorporating various combinations of ABR, SAM EFR, and RAM EFR measures relative to the best-performing single predictor model (RAM_1000_). The root mean square error (RMSE) was computed on the data pooled across 16 and 32 kHz only for mice in the young, aged noise exposed, and aged groups. Markers indicate the change in RMSE of the corresponding model averaged across all repeats and folds, colors indicate the DPOAE adjustment applied, and error bars indicate the standard error of the mean (SEM) of the change in RMSE across all repeats and folds. Dashed line indicates the reference model (RAM_1000_). The ABR_60_ and RAM_110,f0-4_ models are also plotted because they showed additional improvements relative to similar versions of the model included in the main analysis (Figure 6). Asterisks next to the model names indicate combinations that were added after review of the results shown in previous figures. All other comparisons were planned prior to the start of the study.

**Table 7.**
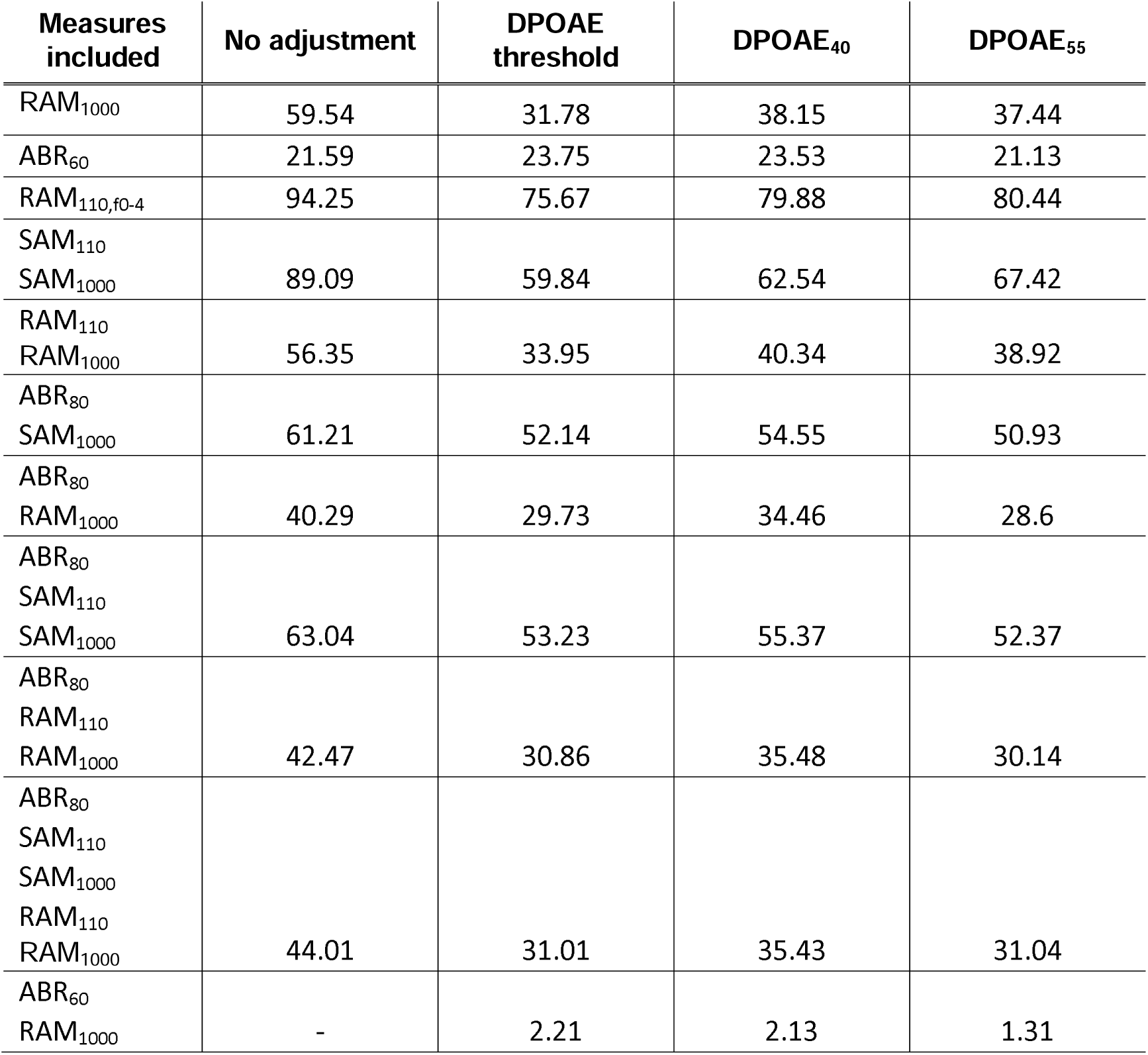
ΔAICc values for combination evoked potential models relative to the best performing model. ΔAICc is computed for each model relative to the best performing model (indicated by a dash). The ABR_60_ and RAM_110,f0-4_ models are shown because they showed additional improvements relative to similar versions of the model included in the main analysis (**Figure 6**).

## Discussion

Adjusting for DPOAEs generally improved the ability of the models to predict synapse numbers, particularly for the SAM EFR models. The differing impact of the DPOAE adjustment on the prediction performance of the SAM versus RAM EFR models is consistent with computational modeling studies suggesting that the sharp onset of the RAM EFR stimulus makes it less vulnerable to impacts from OHC dysfunction than the SAM EFR stimulus (Vasilkov et al., 2021). When assessing models that incorporated ABR wave 1 amplitude, it was not necessary to correct for DPOAEs to yield models that performed well (**Figures 6B, 9 and 10**). This suggests that in human studies of synaptopathy, it may not be necessary to statistically adjust for OHC function when using ABR wave I amplitude as an indicator of synaptopathy.

In mice with no synaptopathy or broad synaptopathy, the RAM_1000_ EFR model was more accurate at predicting synapse numbers than the other single evoked potential models, supporting findings of computational modeling that suggest that the RAM EFR is a better indicator of synaptopathy than the SAM EFR (Vasilkov et al., 2021). The better performance of the RAM_1000_ model as compared to the RAM_110_ model is consistent with animal studies demonstrating that the auditory nerve is the primary generator of the EFR at higher modulation frequencies (Parthasarathy and Kujawa, 2018; Shaheen et al., 2015). Unfortunately, it is unclear if a response to the RAM_1000_ EFR can be obtained in humans. A key limitation when translating this work to humans is that the equivalent rectangular bandwidth depends on cochlear frequency. At carrier frequencies of 16 and 32 kHz, the equivalent rectangular bandwidth in mice is ∼3 and 10 kHz, respectively (May et al., 2006). In humans, comparable carrier frequencies might be 4 and 8 kHz, which translate to equivalent rectangular bandwidths of 0.5 and 1 kHz, respectively (Moore and Glasberg, 1983). Given that the SAM and RAM EFR have sidebands at 𝑓_𝑐_ ± 𝑘 ⋅ 𝑓_𝑚_ where 𝑘 = 1 for SAM and 𝑘 = 1 … ∞ for RAM (**Figure 11**), some stimulus components may fall outside the critical bandwidth of the cochlear filter. For example, for a 4 kHz EFR carrier frequency modulated at 1000 Hz, the sidebands fall outside the equivalent rectangular bandwidth for 4 kHz, while the sidebands for a 16 kHz EFR carrier modulated at 1000 Hz do not fall outside the equivalent rectangular bandwidth for 16 kHz. Despite this caveat, McHaney et al. (2024) demonstrated that the SAM EFR using a 3 kHz carrier with a modulation frequency of 1024 Hz can be obtained in young adults with normal hearing using an ear canal electrode (tiptrode). In gerbil, successful measurement of the RAM_1000_ EFR to a 4 kHz carrier indicates that it is possible to generate a response to this stimulus in an animal with low frequency hearing (**Figure 12**). However, even with these demonstrations of successful measurement of the SAM_1000_ and RAM_1000_ EFR to low frequency carriers, questions regarding the interaction between EFR magnitude and critical bandwidth remain. Thus, future research should investigate the feasibility of collecting the RAM_1000_ EFR in humans and test whether the RAM_1000_ EFR remains a superior predictor of synapse number even in animal models with low frequency hearing.

**Figure 11:**
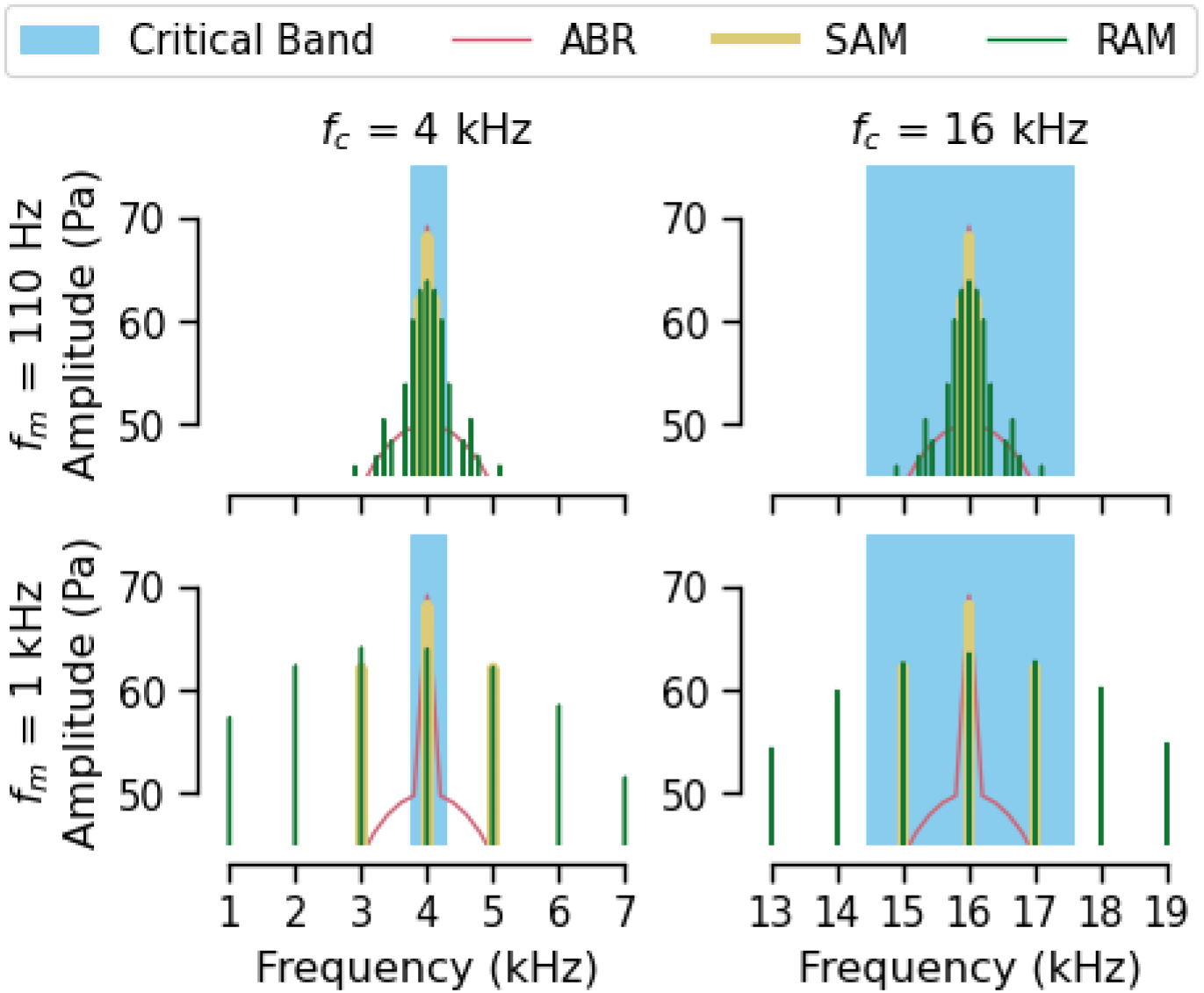
Spectral components of 4 kHz and 16 kHz ABR and EFR stimuli relative to the estimated critical bandwidth. For the RAM and SAM EFR stimuli, the spectral components are produced by modulation of a tonal carrier. Spectral components are positioned at 𝑓_𝑐_ ± 𝑘 ⋅ 𝑓_𝑚_ where 𝑓_𝑐_ is the carrier frequency, 𝑓_𝑚_ is the modulation frequency and *k* is 1 for SAM (yellow) or any number between 1 and ∞ for RAM (green). The shaded blue area indicates the critical bandwidth for 𝑓_𝑐_. For f_c_ = 16 kHz the dominant spectral components fall within the critical band, while for f_c_ = 4kHz they do not. The ABR (pink) is a narrowband stimulus with the dominant power at the stimulus frequency, staying within the critical band for both 4 kHz and 16 kHz stimuli. For comparison, all stimuli are matched in level at 70 dB SPL.

**Figure 12:**
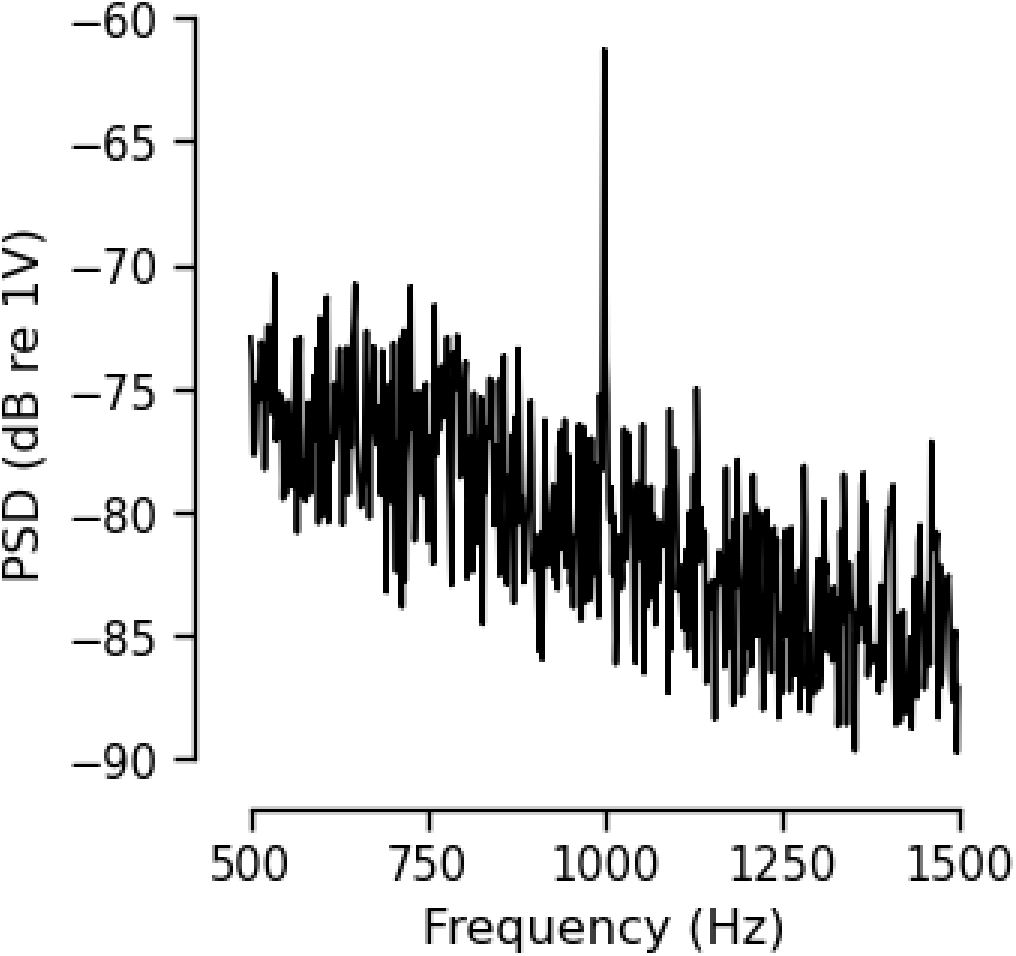
RAM_1000_ EFR obtained in gerbil. Plot shows a recording of the RAM_1000_ EFR in gerbil for a carrier frequency of 4 kHz. A clear peak is observed at the modulation frequency of 1000 Hz.

The synapse prediction performance of the various ABR wave 1 amplitude models indicate that models using lower stimulus levels (40-70 dB SPL) performed better than the model using a high stimulus level (80 dB SPL). Previous human studies of synaptopathy have typically used high ABR stimulus levels (reviewed in Bramhall, 2021), partly to ensure that wave I amplitude can be identified in all participants. Future studies may want to use the lowest ABR stimulus level that still allows for reliable measurement of wave I amplitude even if it requires increasing the number of averages and/or using electrode configurations (i.e., tiptrodes) that improve the signal-to-noise ratio.

Although other EFR processing approaches performed well for some EFR metrics, models using the sum of the absolute power at f_0_-f_4_ were consistently among the best performing of both the SAM and RAM EFR models regardless of modulation frequency. This suggests that future EFR studies should consider using this processing approach. However, there was little difference among the different approaches for the SAM EFR modulated at 110 Hz, suggesting that the results of previous human studies that used different EFR processing approaches would likely not be impacted by the type of processing used.

The improved synapse prediction resulting from combining RAM_1000_ EFR and ABR_60_ wave 1 amplitude in the same model suggests that these measures capture complementary information about synaptopathy. While both the ABR and RAM_1000_ EFR are primarily driven by auditory nerve activity, they may differentially reflect the contributions of spontaneous rate subgroups (Bharadwaj et al., 2014). ABR wave 1 amplitude, measured with an interleaved approach to minimize adaptation, predominantly reflects the initial, onset-driven firing of auditory nerve fibers, a response characteristic of high spontaneous rate fibers. Conversely, the sustained nature of the RAM EFR probes the adapted, steady-state firing and phase-locking abilities of the nerve fibers, a feature more typical of low spontaneous rate fibers. Thus, the ABR and RAM EFR likely provide distinct windows into the functional contributions of different auditory nerve fiber populations.

Of all the evoked potential models, only the ABR wave 1 amplitude model performed better than chance at predicting the number of synapses in mice with focal synaptopathy. However, even the ABR model performed poorly, with prediction errors of approximately four synapses (versus approximately two synapses for the RAM_1000_ model in cases of broad synaptopathy). The poor performance of the ABR wave 1 amplitude model in predicting focal synaptopathy is somewhat surprising given that much of the previous work on the relationship between ABR wave 1 amplitude and synaptopathy was done in animal models of focal noise-induced synaptopathy (e.g., Kujawa and Liberman, 2009). However, this result is consistent with a previous attempt to predict the number of synapses per IHC from ABR wave 1 amplitude and DPOAE levels using a partial least squares regression model (Bramhall et al., 2018), which also overestimated the number of synapses in mice with focal synaptopathy. In contrast to the results of the current study, a previous study of mice with noise-induced synaptopathy showed a linear relationship between mean ABR wave 1 amplitude and mean number of synapses per IHC in groups of mice with temporary threshold shifts and in groups of mice with permanent threshold shifts after adjustment for auditory thresholds (Fernandez et al., 2020). However, the noise exposures in that study resulted in a broader configuration of synaptopathy with a minimum of 20% synaptic loss at 16 kHz, while in the current study neither group of noise exposed mice had synaptic loss at 16 kHz compared to young mice. In this study, the poorer performance of the RAM_1000_ EFR model compared to the ABR model when predicting focal synaptopathy may be due to the broader range of spectral components in the RAM stimulus (**Figure 11**), which will drive regions of the cochlea outside the focal lesion. Further work with a larger dataset of mice with varying degrees of focal synaptopathy may offer insights that can be used to fine-tune evoked potential model predictions for focal synaptopathy.

While the evoked potential models failed to predict focal synaptopathy in the current study, this may not preclude diagnosis of human synaptopathy. Synapse counts from human temporal bones do not show any evidence of focal, or frequency-specific, synaptopathy (Wu et al., 2019; Wu et al., 2021). Even if focal synaptopathy does occur immediately following noise exposure in humans, it likely expands into broad synaptopathy over long time periods similar to what is observed in mice that are noise exposed and then aged (**Figure 2E**; Fernandez et al., 2015).

In conclusion, in mice with no synaptopathy or broad synaptopathy, the RAM_1000_ EFR model adjusted for DPOAEs resulted in the best prediction of synapse numbers out of all the single evoked potential models, while a model that combined the RAM_1000_ EFR and ABR_60_ wave 1 amplitude without a DPOAE adjustment provided the best overall performance out of all the predictive models that were evaluated. This suggests that future studies of synaptopathy should measure the RAM_1000_ EFR, ABR wave 1 amplitude, and DPOAEs. However, concerns regarding the interaction between the critical bandwidth and the spectral components of the EFR stimuli require further study to establish whether the RAM_1000_ EFR remains a strong predictor of cochlear synapse number even at the low carrier frequencies needed to obtain this measure in humans. If the RAM_1000_ EFR turns out not to be a viable metric for use in humans, the results of this study suggest that ABR wave I amplitude, particularly for lower stimulus levels, is also strong measure of synaptopathy. None of the evoked potential models performed well at predicting focal synaptopathy, although the ABR wave 1 amplitude model performed the best. Additional work will be necessary to identify physiological measures that can accurately predict focal synaptopathy.

